# Rapid transgenerational adaptation in response to intercropping increases facilitation and reduces competition

**DOI:** 10.1101/2022.01.14.476288

**Authors:** Laura Stefan, Nadine Engbersen, Christian Schöb

## Abstract

By capitalising on positive biodiversity-productivity relationships, intercropping provides opportunities to improve agricultural sustainability^1^. However, intercropping is generally implemented using commercial seeds that were bred for maximal productivity in monocultures, which might limit the benefits of crop diversity on yield^2,3^. Plants can adapt over generations to the level of surrounding plant diversity, notably through increases in niche differentiation^4^. However, this adaptation potential and the corresponding yield benefit potential have not been explored in annual crop systems. Here we show that plant–plant interactions among annual crops evolved towards increased facilitation and reduced competition when the plants’ coexistence history matched their current diversity setting, which led to an increase in overyielding of up to 58%. These higher yield benefits were linked to character convergence between species sharing the same coexistence history for two generations. Notably, the six crop species tested converged towards taller phenotypes with lower leaf dry matter content when grown in mixtures. This study provides the first empirical evidence for the importance of parental diversity affecting plant–plant interactions and ecosystem functioning of the following generations in annual cropping systems. These results have important implications for diversified agriculture as they demonstrate the yield potential of targeted cultivars for intercropping, which can be achieved through specific breeding for mixtures.

---

Following decades of studies demonstrating the positive relationship between species diversity and plant primary productivity in natural systems^5,6^, intercropping, i.e. growing more than two species in the same field during the same period, has been increasingly considered as a promising option to increase agricultural sustainability^1,7^. The productivity benefits of increasing species diversity rely on two main mechanisms, namely selection effects and complementarity effects, the latter encompassing both facilitation and niche differentiation^8,9^. In perennial natural grasslands, complementarity effects have been shown to increase over time due to evolutionary processes^4,10,11^. Notably, greater species complementarity can result from evolutionary changes^12^ – i.e. changes in gene frequency – or from heritable epigenetic changes^13^ affecting species traits in response to surrounding plant diversity, which either increases niche differentiation (i.e. reduces competition) or facilitation^14^. The evolutionary potential of plant–plant interactions in diverse communities has tremendous implications for the diversification of agricultural systems^15^. This is of particular relevance for mixed cropping systems, where the use of commercial seeds domesticated and bred for maximum yield in monoculture is the norm, which may compromise the diversity benefits^2,3,16–18^. Despite the paramount importance of this question, the yield potential of mixture-adapted varieties is, to our knowledge, unknown, as are the character differences of monoculture-compared to mixture-adapted crops. Therefore, in this project, we determined whether and how crop species adapt over three generations to the level of plant diversity that they are surrounded by. We investigated how plant–plant interactions, i.e. competition and facilitation, and plant traits changed and evolved within different coexistence histories over time, and whether these changes translated into yield benefits. To that end, we conducted an intercropping experiment in Switzerland with six different crop species commonly cultivated in Europe and belonging to four functionally different phylogenetic groups. The mesocosms included monocultures, 13 different 2-species mixtures, four different 4-species mixtures, and isolated single plants, and was replicated in two different fertilizing conditions. We selected open-pollinated varieties as seed source to provide the genetic variability needed for evolutionary processes to occur. To assess potential transgenerational changes, we repeated the experiment over the course of three years with seeds from plants grown from either monocultures, mixtures, or single individual plants of the previous year (Fig. 1, Fig. 5).

**Figure 1.**
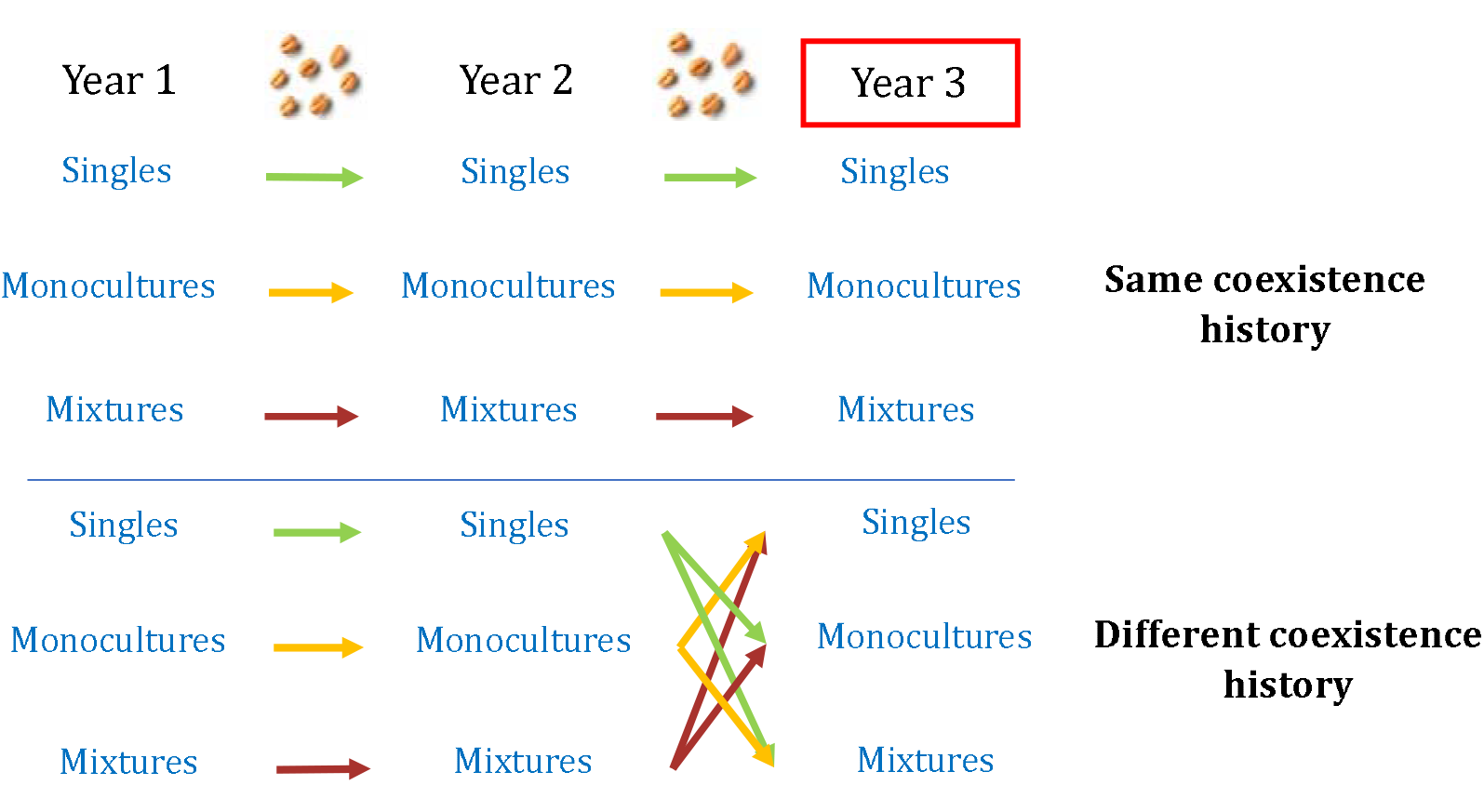
Experimental design. Six crop species were used to sow single plant individuals (6), monocultures (6), 2-species mixtures (13) and 4-species mixtures (4) in 2018 (Year 1); seeds were collected at the end of the growing season and resown in 2019 (Year 2) in the same diversity setting as their previous generation. Seeds were collected again and resown in 2020 (Year 3), this time either in the same community their seeds were collected from [same coexistence history], or in a community different to the one of their parents [different coexistence history] (*n* = 468 plots). This process was replicated in two different fertilizing conditions. We expected that crops growing in the same community as their parents would have adapted over the two generations, and therefore would exhibit less competition and have higher productivity than crops growing in a community different to the one of their parents.

Results from the third year showed that plant–plant interactions shifted towards stronger facilitation and weaker competition when the plants were growing in the same community conditions than their two previous generations (Fig. 2, Extended Data Fig. 1, Extended Data Table 1). More precisely, net interaction index, as well as competition and facilitation indexes, were significantly higher when the crops were grown in the community their seeds were collected from than when they were growing in a community different to the one of their parents (Fig. 2, Extended Data Fig. 1; +54% for the net index, +9% for the competition index, +93% for the facilitation index). Pairwise comparisons further showed that this effect of coexistence history was particularly true in mixtures and only a trend in monocultures, for both fertilizing conditions (Extended Data Table 2). This notably demonstrates that in mixtures, mixture-adapted communities (i.e. with the same coexistence history) exhibited more facilitation and less competition than monoculture-adapted communities or single plant-adapted communities (i.e. with a different coexistence history).

**Figure 2:**
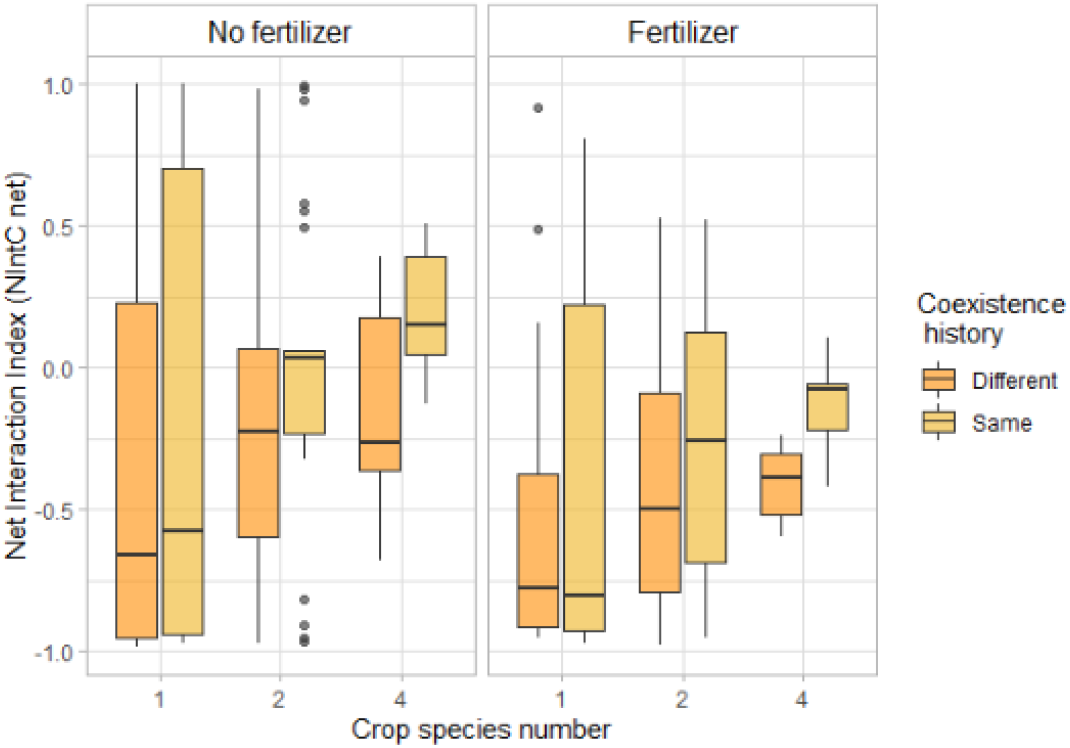
Plant interaction index in response to coexistence history. Net interaction index of monocultures, 2- and 4-species mixtures in response to coexistence history, for fertilized and unfertilized conditions. n =276. This index compares the performance of plants growing in communities to the performance of single plants (see Methods). Negative interaction index indicates competition within a community, positive interaction index indicates facilitation. The closer this index gets to 1, respectively −1, the stronger the facilitation, respectively competition. “Same coexistence history” indicates that crops were grown in the community their seeds were collected from. “Different coexistence history” refers to crops grown in a community different to the one of their parents. The effect of fertilization and coexistence history were highly significant. See Extended Data Table 1 for the complete statistical analysis, and Extended Data Fig. 1 for competition and facilitation indexes. Horizontal lines represent the median of the data, boxes represent the lower and upper quartiles (25% and 75%), with vertical lines extending from the hinge of the box to the smallest and largest values, no further than 1.5 * the interquartile range. Data beyond the end of the whiskers are outlying and plotted individually.

This shift in plant–plant interactions was accompanied by a similar shift in net biodiversity effect (NE) in fertilized plots (Fig. 3a). Net biodiversity effect was calculated following the method of Loreau & Hector (2001) and represents the deviation from the expected yield in the mixture, based on the yield of the corresponding monocultures^8^. We observed that under fertilized conditions, NE was on average 58% higher with the same coexistence history than with a different coexistence history (Fig. 3a, Extended Data Table 3), which corresponded to an increase in total yield ranging from 8 to 22% in mixtures (Fig. 3b). This indicates that in fertilized plots, the yield benefits of crop mixtures were higher with mixture-adapted individuals compared to monoculture-adapted and single-adapted individuals. Interestingly, in unfertilized plots we did not observe the same trend. When looking at the partitioning of net effects into complementarity and selection effects^8^, we only observed a significant effect of coexistence history on selection effects under fertilized conditions for 4-species mixtures (Extended Data Fig. 2b).

**Figure 3:**
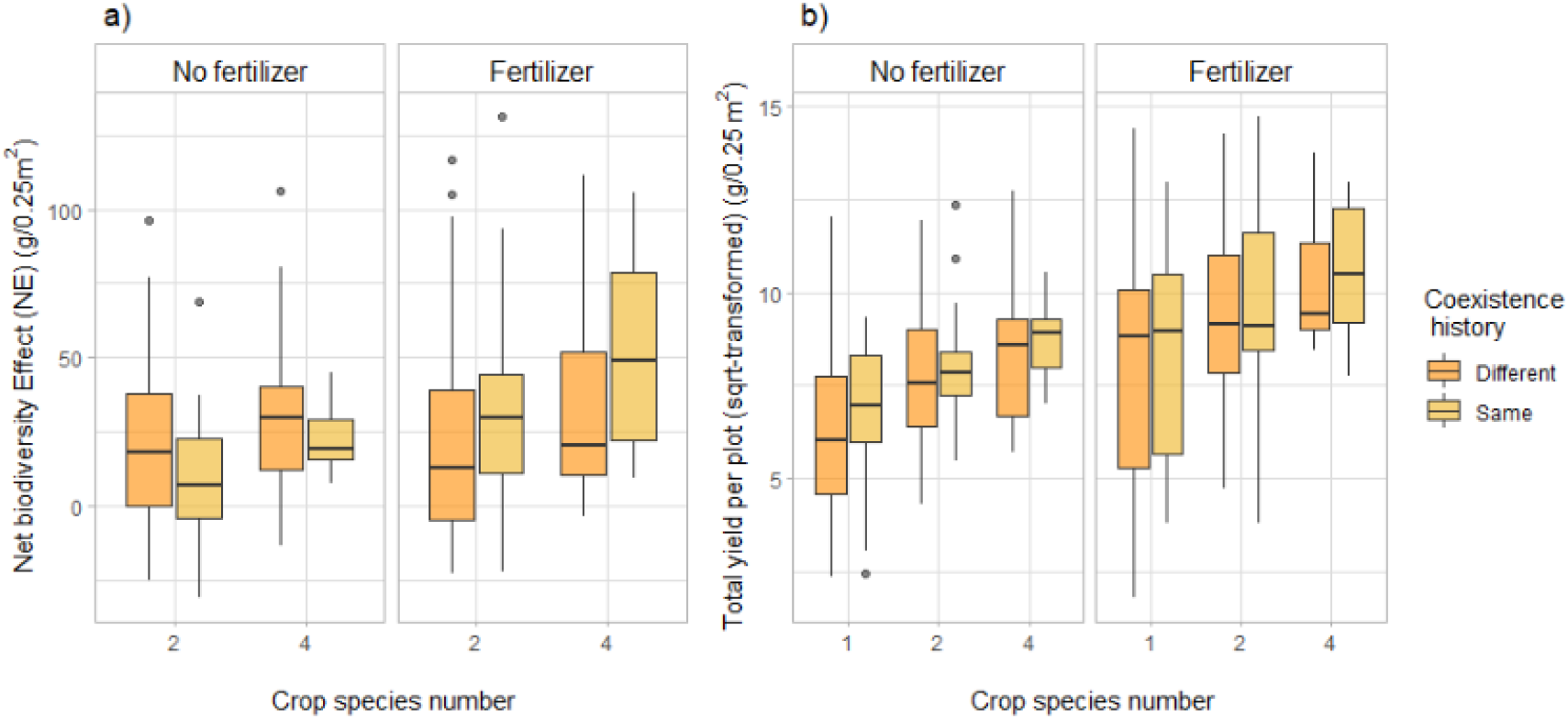
Effects of coexistence history on net biodiversity effects (a) and total yield per plot (b) Effects of coexistence history and crop species number on (a) net biodiversity effect – reflecting the yield advantage of mixtures compared to monocultures – and (b) total yield per plot (square-root transformed) in fertilized and unfertilized plots. (a) n =276; (b) n=204. “Same coexistence history” indicates that crops were grown in the community their seeds were collected from. “Different coexistence history” refers to crops grown in a community different to the one of their parents. See Extended Data Table 3 & 4 for the complete statistical analysis, and Extended Data Fig. 2 for complementarity and selection effects. Horizontal lines represent the median of the data, boxes represent the lower and upper quartiles (25% and 75%), with vertical lines extending from the hinge of the box to the smallest and largest values, no further than 1.5 * the interquartile range. Data beyond the end of the whiskers are outlying and plotted individually.

To investigate the ecological mechanisms behind the shift in plant–plant interactions and biodiversity effects with coexistence history, we measured standard above-ground plant traits and compared the average values as well as coefficients of variation at the species and community levels of single-, monoculture- and mixture-adapted varieties. Following traditional niche theory, we expected that the observed reduction in competition would be linked to an increase in functional trait variation, thereby reflecting an increase in niche differentiation. Surprisingly, we did not observe character displacement – i.e. increased trait variation^4^ – in our intercrop systems, but rather character convergence – i.e. reduced trait variation (Fig. 4). More specifically, we found a reduction in trait variation at the community level, notably of height and leaf dry matter content: the coefficient of variation of height was lower in the same coexistence history treatment compared to a different coexistence history (−9%) (Fig. 4d), and for leaf dry matter content it was 15% lower with the same history compared to a different history (Fig. 4e). Furthermore, the coefficient of variation of mass per seed was also lower under the same history compared to a different history, but this effect was only significant in monocultures (−33%) (Fig. 4f). The community-weighted means of plant traits (CWM, calculated at the community level) further suggest that when growing in the same coexistence history, plants seem to converge towards taller individuals with lower leaf dry matter content. Indeed, the community-weighted mean of leaf dry matter content was significantly lower with the same coexistence history compared to a different history (−3%, Fig 4); height community-weighted mean was – although non-significantly – higher in the case of the same history compared to different coexistence history (Fig 4a). We observed similar responses of height and leaf dry matter content at the species level (Extended Data Fig. 5 & 6, Extended Data Tables 5–9).

**Figure 4:**
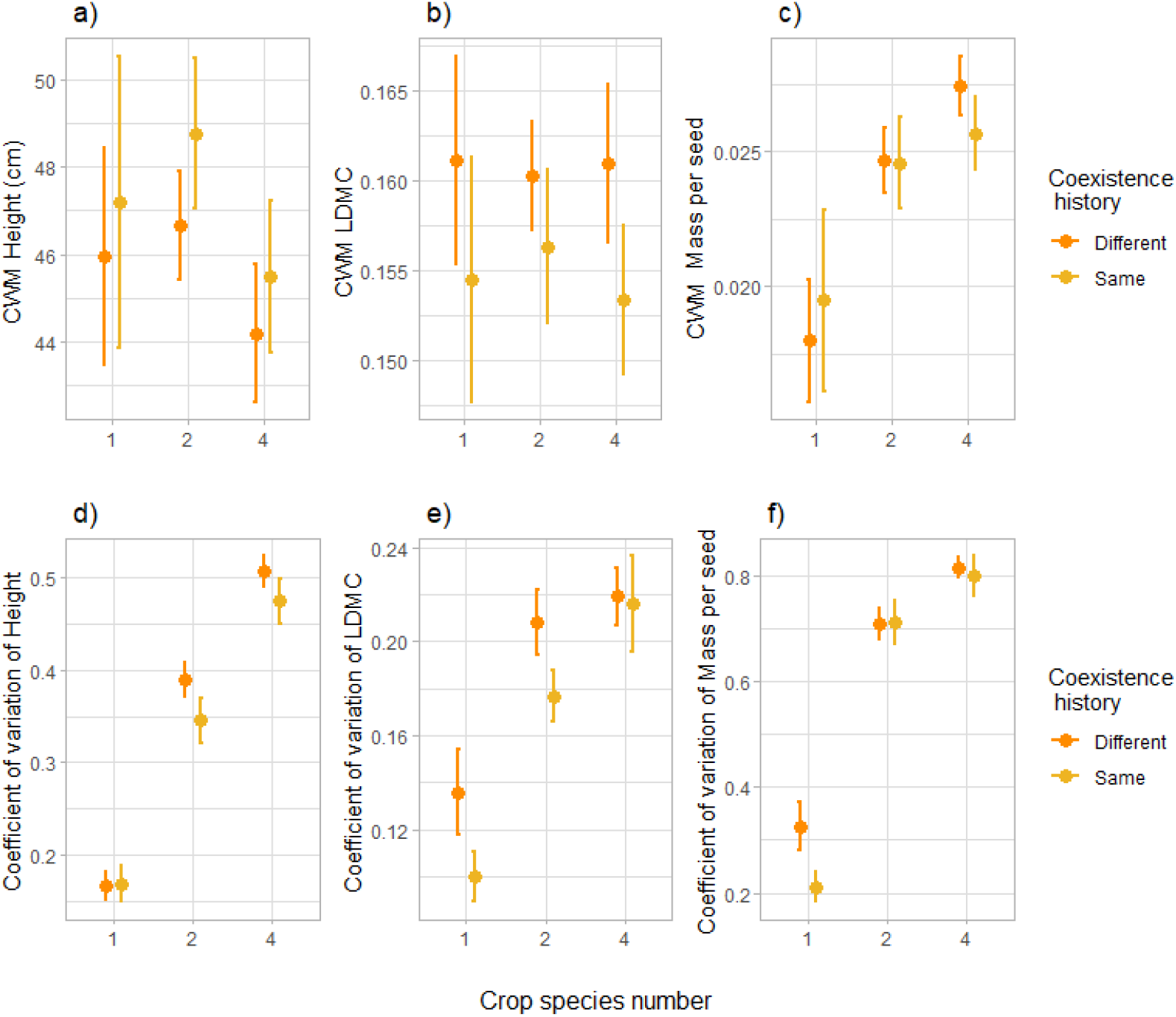
Plot-level traits response to coexistence history. Effects of coexistence history and crop species number on community-weighted mean (CWM) of height (in cm) (a), Leaf Dry Matter Content (LDMC) (b), and mass per seed (in g) (c), and on coefficient of variation at the community level of height (d), Leaf Dry Matter Content LDMC (e), and mass per seed (f). n= 271. “Same coexistence history” indicates that crops were grown in the community their seeds were collected from. “Different coexistence history” refers to crops grown in a community different to the one of their parents. Dots represent the mean values across plots; lines represent the standard error. See Extended Data Tables 10–14 for the complete statistical analysis, and Extended Data Fig. 5–6 as well as Extended Data Tables 5–9 for responses at the species level.

Our research demonstrates that, after only two generations, annual crop plants growing in the same diversity setting as their preceding generations showed reduced competition and increased facilitation compared to plants growing in a different diversity setting as their parents, which led to increased overyielding. We further investigated whether character displacement was responsible for this evolution of plant–plant interactions; contrary to our hypotheses, results did not show evidence for character displacement, but rather for character convergence in plant aboveground traits.

The observed shift in plant–plant interactions are consistent with several grassland studies investigating the effects of community evolution on community productivity and niche differentiation, where it was found that common rapid evolution in plant communities can lead to increases in ecosystem functioning^4,10,11,19^. We indeed observed a positive effect of common community history on the net biodiversity effect (i.e. overyielding), which means that the yield benefit of mixtures compared to monocultures was higher when the plants had been adapted to growing in mixtures (Fig. 3). This can explain why diversity effects generally increase over time^20,21^. Here we did not observe a significant increase in complementarity effect in response to common community history (Extended Data Fig. 2a). However, we observed a similar trend – although nonsignificant – for CE as for NE (Extended Data Fig. 2a): in fertilized plots, CE tended to be higher in the case of a mixture coexistence history, notably in 2-species mixtures. We suggest that the limited timeframe of this study – two generations – might be the reason for the lack of more significant changes in CE and emphasizes the need for longer-term research to confirm or infirm this trend. Surprisingly, selection effects also increased in 4-species mixtures in response to coexistence history (Extended Data Fig. 2b). This is unexpected, as selection effects have not, to our knowledge, been shown to increase over time^22^. However, it might be that this short common community history has favoured a specific species or a specific trait that was particularly plastic or beneficial for fitness^23,24^.

The above-mentioned increases in biodiversity, complementarity and selection effects were only present in fertilized conditions, which could indicate that the benefits of common community history might be dependent on the abiotic conditions. This is nonetheless consistent with several recent studies demonstrating that biodiversity effects are higher in high-inputs systems^2,25,26^, and emphasizes the role of fertilization in driving these effects. Indeed, by promoting crop growth and, consequently, higher competition between plants, fertilization may foster higher benefits of niche differentiation^27–29^.

Overall, increases in biodiversity effects are associated with changes in species traits in response to surrounding plant diversity^4,14,30^. Traditional hypotheses of trait and niche theory indeed predict that when several species co-occur closely together, selection over generations would favour character displacement that would reduce resource overlap and consequently increase niche differentiation^31,32^. Surprisingly, here we found the reverse and observed that a reduction in trait variation favoured increased yield benefits in mixtures. Furthermore, functional diversity – calculated as the volume occupied in the space of the considered traits in this study^33^ – did not respond to common coexistence history (Extended Data Fig. 7 & Table 15). While surprising, this result is not unheard of^23^, and suggests that our plants might have adapted to express the phenotype that would maximise their fitness^34–36^. This ideal phenotype is, in our mixture communities, characterized by taller plants with lower leaf dry matter content, the latter indicating soft leaves associated with rapid biomass production^37^, and consequently less resource-conservative strategies^38^. Lower leaf dry matter content has recently been associated with lower parental or ambient competition^39^, which is consistent with our results of plant–plant interaction intensities. The traits examined here did not allow to understand the mechanisms behind the observed reduction in competition; we suggest that other traits or processes not measured in this experiment might have responded to the coexistence history treatment. Notably, there could be a shift in below-ground traits, such as root-associated traits^39^, or temporal differentiation of resource capture ^40^, such as light. We indeed observed a significant increase in light capture ability in plants coming from the same diversity setting compared to the same communities but a different coexistence history (Extended Data Fig. 8 & Table 16), which indicates that plants used to growing in the same diversity setting during several generations might capture the resources more fully than plants coming from a different diversity setting. However, here we rely on our light interception measurements and suggest more longer-term studies to understand changes in the use of other resources, such as nutrients or water, and how this is associated to plant traits. Furthermore, the scope of this study did not allow us to investigate the mechanisms behind these changes in plant-plant interactions and traits in response to coexistence history. The adaptation response might be genetically-based and due to natural selection^11^, as we specifically selected open-pollinated varieties in order to ensure a minimum amount of genetic variability. Furthermore, outcrossing could have occurred in the first year of this experiment, as we had a similar experiment running in the same experimental garden with Spanish varieties from the same species^27,41^. However, considering the short timeframe of this study and the low rate of outcrossing in most of our species, epigenetic changes – i,e, stable heritable changes in cytosine methylation – might also have played an important role as potential evolutionary mechanisms^13,42–46^.

For the first time, our study provides empirical evidence for rapid transgenerational adaptation in response to diversity history in annual crop communities. Notably, we demonstrated that when plants were coming from the same diversity setting as their parents, plant–plant interactions shifted towards reduced competition and increased facilitation. This effect was particularly true for mixtures and translated into enhanced overyielding under fertilized conditions. This reduction in competition was surprisingly not linked to character displacement, but we instead observed character convergence towards taller plants with lower leaf dry matter content. This research emphasizes the importance of considering transgenerational effects of diversity for crop mixtures. This is particularly relevant for breeding programs and highlights the need of including diversity when breeding for crop mixtures, in order to design varieties specifically adapted for intercropping^17^.

## Methods

### Study sites

The Crop Diversity Experiment took place in 2018, 2019, and 2020 in an outdoor experimental garden located at the Irchel campus of the University of Zurich, Switzerland (47.3961 N, 8.5510 E, 508 m a.s.l). Zurich is characterized by a temperate climate^27^. The experimental garden was irrigated during the growing season with the aim of maintaining a sufficient amount of water for optimal plant growth. The dry threshold of soil moisture was set at 50% of field capacity, with a target soil moisture of 90% of field capacity. Whenever dry thresholds were reached (measured through PlantCare soil moisture sensors (PlantCare Ltd., Switzerland), irrigation was initiated, and water added until reaching the target value.

Each experimental garden consisted of square plots of 0.25 m^2^. The uppermost 30 cm were filled with standard, not enriched, agricultural soil coming from the local region. This soil consisted of 45 % sand, 45 % silt, and 10 % clay, and initially contained 0.19 % nitrogen (N), 3.39 % carbon (C), and 332 mg total phosphorous (P)/kg, with a mean pH of 7.25. Beneath that, there was local soil of uncharacterized properties that allowed unlimited root growth. The plots were embedded into larger beds of 7 × 1 m, each bed containing 28 plots. Inside a bed, plots were separated from each other by metal frames. While the relatively small plot sizes allowed us to undertake a large experiment under environmentally highly controlled but realistic outdoor conditions, some variables can suffer edge effects and interferences with neighbouring plots. However, such effects would probably increase residual variation more than between-treatment variation, because randomization was used to prevent confounding of between-plot interactions with treatments. In the only relevant study of which we are aware, the biodiversity–productivity relationship in herbaceous communities was not affected by plot size^47^ while a recent theoretical study showed that, if anything, biodiversity effects should increase with plot size^48^. We therefore assume that effect size in our experiment, if anything, is probably rather conservatively estimated compared with that in studies using larger plot sizes.

Every year, we fertilized half of the beds with N, P and potassium (K) at the concentration of 120 kg/ha N, 205 kg/ha P, and 120 kg/ha K. Fertilizers were applied three times per year, namely once just before sowing (50 kg/ha N, 85 kg/ha P, 50 kg/ha K), once when wheat was at the tillering stage (50 kg/ha N, 85 kg/ha P, 50 kg/ha K), and once when wheat was flowering (20 kg/ha N, 34 kg/ha P, 20 kg/ha K). The other half of the beds served as unfertilized controls. In 2018, we randomly allocated individual beds to a fertilized or non-fertilized control treatment. In the following years, we kept the initial fertilization treatment allocation.

### Crop species

Experimental communities were constructed with six annual crop species of agricultural interest. We selected only seed crops with similar growth requirements in terms of climate and length of growing season, and with similar plant sizes to fit at least 40 individuals in the rather small plots. The six species belong to four different phylogenetic groups with varying functional characteristics: we first separated monocots [*Triticum aestivum* (wheat, C3 grass, Poaceae) and *Avena sativa* (oat, C3 grass, Poaceae)] and dicots. Among the dicots, we differentiated between suparasterids [*Coriandrum sativum* (coriander, herb, Apiaceae)] and superrosids. Among the superrosids, we separated legumes [*Lens culinaris* (lentil, legume, Fabaceae)] from non-legumes [*Linum usitatissimum* (flax, herb, Lineceae) and *Camelina sativa* (false flax, herb, Brassicaceae)]. Furthermore, we chose crop varieties that were locally adapted and commercially available in Switzerland (Table 1).

**Table 1.**
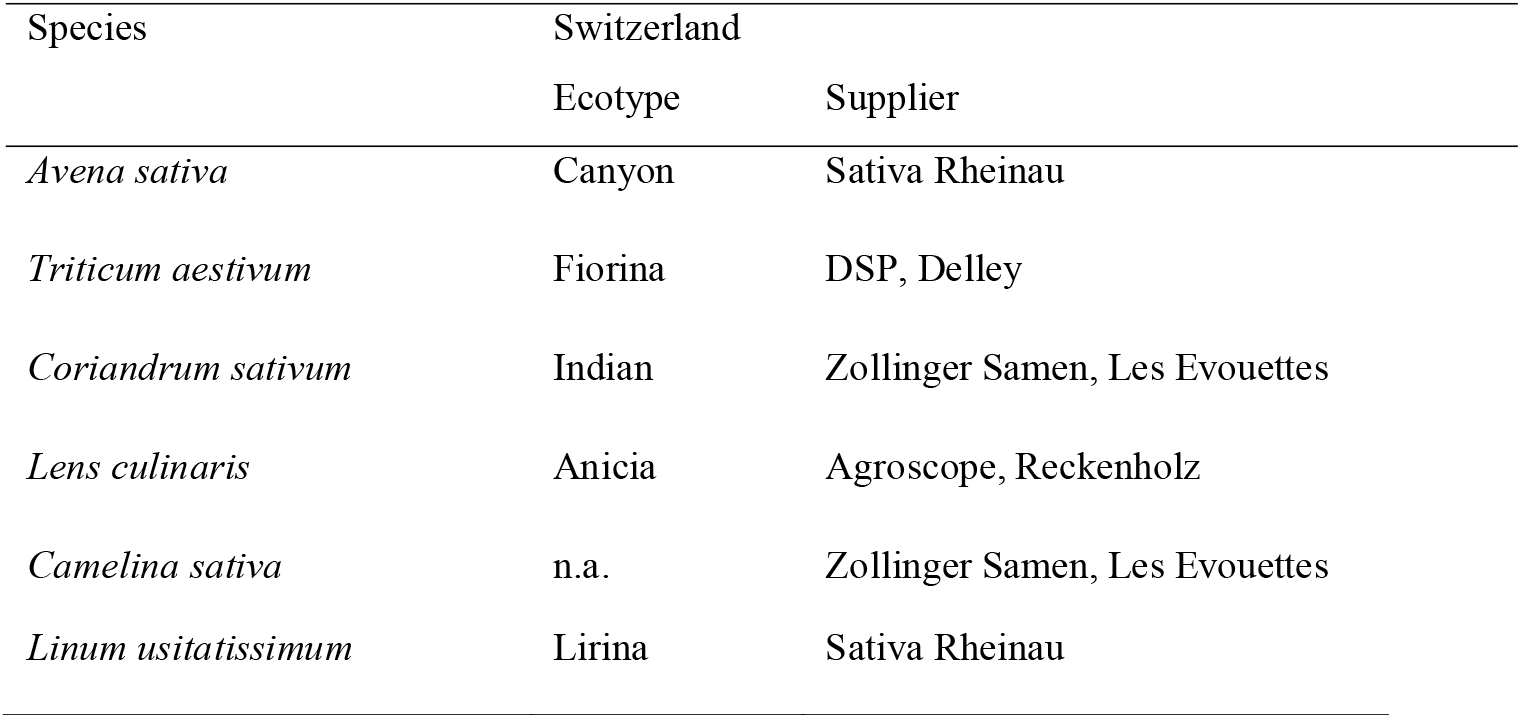
List of crop species ecotypes and their suppliers.

*Avena Sativa* (oat) is mainly self-pollinating, with outcrossing rates of around 1%^52^. The variety Canyon was acquired in 2014 through conventional selection processes.

*Triticum aestivum* (wheat) is principally self-pollinating, with outcrossing rates generally between 1 and 4%^49,50^, although some cultivars have been shown to have outcrossing rates up to 8%^51^. Fiorina is an accession originating from Switzerland, acquired in 2015, specifically for organic agriculture.

*Coriandrum sativum* (coriander) has a generally high genetic variability, with studies showing up to 70.46% polymorphism, indicating the presence of high degree of molecular variation in the studied coriander varieties^58,59^. The variety that we used originally came from an Indian market and was not a fixed variety, which ensured a minimum of genetic variability. The flowers of coriander are self-incompatible but plants are self-compatible. Geitonogamy is therefore common. Cross-pollination is facultative but can reach up to 20%^60^. *Lens culinaris* (lentil) is mainly self-pollinating; depending on the cultivar, outcrossing rates reach between 1 and 5%^53^.

*Camelina sativa* (camelina) is mainly self-pollinating, with outcrossing rates of less than 1%^61,62^. In the study, we used a local landrace that was not a fixed variety.

*Linum usitatissimum* (flax) is mainly self-pollinating but outcrossing does occur, at a rate of 1-5%^54^. Lirina, the variety of Linum that we used has been defined by ProSpecieRara as a rare or ancient variety. ProSpecieRara ensures the preservation of rare traditional varieties^55^. Furthermore, studies have shown that linseed varieties have higher genetic variability than fiber flax and should therefore be considered as valuable genetic resources^56,57^.

### Experimental crop communities

Experimental communities consisted of single plots with one individual, monocultures, 2- and 4-species mixtures (Fig. 5). We planted every possible combination of 2-species mixtures with two species from different phylogenetic groups and every possible 4-species mixture with a species from each of the four different phylogenetic groups present (Table 2). We replicated the experiment two times with the exact same species composition, except for single individuals which were replicated 4 times. Monoculture and mixture plots were randomized among plots and beds within each fertilizer treatment, while single plant plots were randomly allocated to plots in separate beds in order to minimize interference among neighbouring plots. Each monoculture and mixture community consisted of one, two or four species planted in four rows. Two species mixtures were organized following a speciesA|speciesB|speciesA|speciesB pattern. The order of the species was chosen randomly. For 4-species mixtures, the order of the species was also randomized. Density of sowing differed among species groups and was based on current cultivation practices: 160 seeds/m2 for legumes, 240 seeds/m2 for superasterids, 400 seeds/m2 for cereals, and 592 seeds/m2 for superrosids. Each year, seeds were sown by hand in early April.

**Figure 5:**
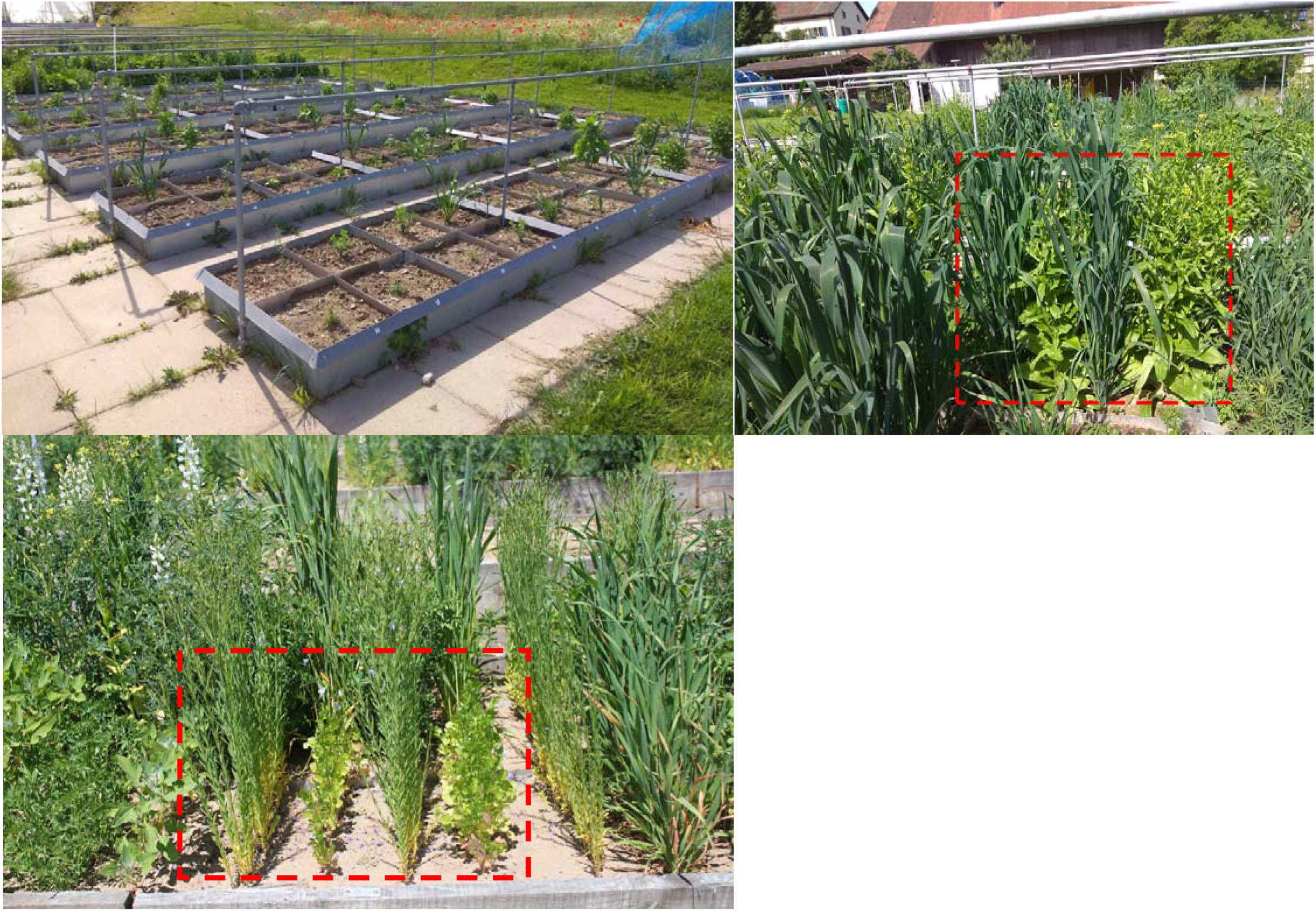
Pictures of the experimental plots. Top-left: part of the experimental garden, showing the plots within beds, and planted with single individuals. Top-right: a plot is outlined in red, showing a 2-species mixtures, with a cereal (wheat or oat) alternated with camelina. Bottom-left: a plot is outlined in red, showing a 2-species mixtures, with flax alternated with coriander.

**Table 2.** List of species mixture combinations.

| *Monoculture* | *2-species mixtures* | *2-species mixtures* | *4-species mixtures* |
| --- | --- | --- | --- |
| Avena | Avena-Lens | Lens-Linum | Avena-Lens-Linum-Coriandrum |
| Triticum | Avena-Linum | Lens-Camelina | Avena-Lens-Camelina-Coriandrum |
| Lens | Avena-Camelina | Lens-Coriandrum | Triticum-Lens-Linum-Coriandrum |
| Linum | Avena-Coriandrum | Linum-Coriandrum | Triticum-Lens-Camelina-Coriandrum |
| Camelina | Triticum-Lens | Camelina-Coriandrum |  |
| Coriandrum | Triticum-Linum | Triticum-Coriandrum |  |
|  | Triticum-Camelina |  |  |

### Adaptation treatment

In 2019, we used the seeds collected in 2018 to add a coexistence history treatment: we repeated the experiment with seeds coming from single individuals, monocultures, and mixtures, respectively. This means that each plot described above was repeated three times: once with seeds coming from single plants, once with seeds coming from monoculture plants, and once with seeds coming from mixture plants. We respected the fertilizing treatment, i.e. there was a history treatment for each fertilizing condition. When planting the mixtures with a mixture history, we specifically used seeds coming from the same species combination. When planting the monocultures and singles with a mixture history, we used seeds coming from a common pool combining all 4-species mixtures.

In 2020, we repeated this process and selected seeds from 2019 to sow the single and community plots. We only selected seeds that had a “pure” history, i.e. that were always grown in the same coexistence history (for instance, for single history seeds in 2020 we selected only seeds that were grown as singles also in 2018 and 2019).

### Data collection

#### Photosynthetically Active Radiation (PAR)

Interception of PAR by the plant canopy was measured weekly with a LI-1500 (LI-COR Biosciences GmbH, Germany). In each plot, three PAR measurements were taken around noon by placing the sensor on the soil surface in the center of each of the three in-between rows. Light measurements beneath the canopy were compared to ambient radiation through simultaneous PAR measurements of a calibration sensor, which was mounted on a vertical post at 2 m above ground in the middle of the experimental garden. FPAR (%) indicates the percentage of PAR that was intercepted by the crop canopy.

#### Traits measurements

At the time of flowering, three individuals per crop species per plot were randomly marked. We measured the height of each individual with a ruler from the soil surface to the highest photosynthetically active tissue. We then measured plant width with a ruler by taking the largest horizontal distance between two photosynthetically active tissues. We sampled one healthy leaf from each marked individual and immediately wrapped this leaf in moist cotton; this was stored overnight at room temperature in open plastic bags. The following day, we removed any excess surface water on the leaf and weighed it to obtain its water saturated weight ^63^. Then this leaf was scanned with a flatbed scanner (CanoScan LiDE 120, Canon), oven-dried in a paper envelope at 80°C for 72 hours, and subsequently reweighed to obtain its dry weight. We calculated Leaf Dry Matter Content (LDMC) as the ratio of leaf dry mass (g) to water saturated leaf mass (g). Using the leaf scans, we measured leaf area with the image processing software ImageJ ^64^. Specific Leaf Area (SLA) was then calculated as the ratio of leaf area (cm2) to dry mass (g).

#### Plot grain yield and biomass

Grain yield and aboveground biomass of each crop species was determined per plot at maturity. This corresponded to July/August. As time of maturity slightly varied among the different crop species, we conducted harvest species by species. We clipped plants right above the soil surface and separated seeds from the vegetative parts. Seeds were sun-dried for five days and weighed. Biomass was oven-dried at 80 °C until constant weight and weighed.

#### Individual yield and biomass

We harvested the three marked individuals for the trait measurements separately; we separated seeds from aboveground biomass and they were both dried and weighed as previously mentioned. Furthermore, for each marked individual we weighed ten randomly selected seeds to obtain the mass per seed.

### Data analyses

All analyses were performed using R version 4.1.0^65^.

#### Plant Interaction Index

Plant interaction intensity in the plots was calculated for each marked individual by means of the neighbor-effect intensity index with commutative symmetry NIntC^66^:

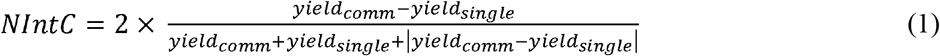

where *yield*_*single*_ is the yield of a single plant grown in isolation, and *yield*_*comm*_ is the yield of an individual of the same species when grown in a community. NIntC values of all species (*a,b,c,d*) composing the community (i.e. species a in case of a monoculture and species a to d in case of a mixture of four species) were averaged and subsequently weighted by their proportional abundance 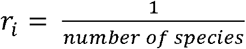 to calculate the mean net interaction in the community (NIntCnet):

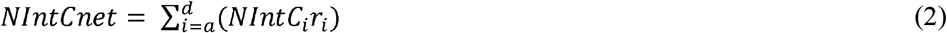

We then partitioned this net interaction index into its facilitation and competition components: NIntC facilitation was obtained subsetting those individuals with a positive NIntC value (i.e. with increased performance in communities compared to single plant individuals without neighbour interactions) and calculating the mean of all the species per plot weighed by their relative abundance; NIntC competition was obtained by subsetting those individuals with a negative NIntC value (i.e. with reduced performance in communities compared to single plant individuals without neighbour interactions) and calculating the mean of all the species per plot weighed by their relative abundance.

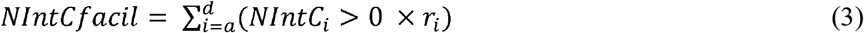

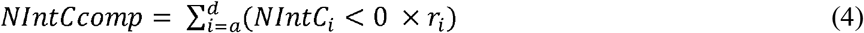

#### Net biodiversity effect

For all mixture communities we quantified the net biodiversity effect (NE) and its two components, the complementarity and selection effects according to Loreau and Hector^8^:

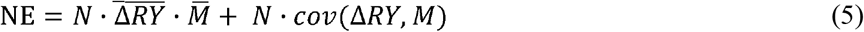

where N is the number of species in the plot, ΔRY is the deviation from expected relative yield of the species in mixture in the respective plot, which is calculated as the ratio of observed relative yield of the species in mixture to the yield of the species in monoculture, and M is the yield of the species in monoculture. The first component of the biodiversity effect equation 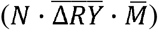 is the complementarity effect (CE), while the second component (*N· cov(ΔRy, M*)) is the selection effect (SE).

#### Total crop yield

To assess crop performance, we calculated total crop yield per plot as the sum of total seed mass per species.

#### Trait analyses

Traits were analysed both at the species-level and at the plot-level. At the species level, we calculated the mean and coefficient of variation (CV) per species for each trait per plot. At the plot-level, we calculated Community-Weighted-Means (CMW) based on biomass per species, and coefficient of variation per plot for each trait.

Functional richness (FRic) was calculated in each plot using the function *dbFD* from the package *FD*^67^, by measuring the convex hull volume occupied by the individuals of a plot in the space of the considered traits.

To analyze the effects of the experimental treatments on NIntCnet, NIntCfacil, NIntCcomp, NE, CE, LER, total crop yield, FRic, and CWM and CV per plot, we used generalized linear mixed models using the function *lmer*. Fixed factors included fertilizing condition (yes or no), coexistence history (considered as “same” or “different”), crop species number (2 vs 4) nested in monoculture vs mixture, as well as the interactions between them. Species composition and bed were set as random factors. Effect sizes were calculated from marginal means obtained using the function *emmeans,* and pairwise comparisons were calculated using Tukey tests from the *emmeans* function^68^. To analyze the effects of the experimental treatments on the mean and coefficient of variation of the different traits per species (height, width, SLA, LDMC, mass per seed, respectively), we used generalized linear mixed models using *lmer* with the same fixed factors as previously described. Species, species composition and bed were set as random factors. The response variables were log-transformed or square-root-transformed where needed. To analyse the response of FPAR, we used similar linear mixed models as described above, but added day of year as a random factor. For all models, we tested for normality of the residuals using a Shapiro-Wilk test and homogeneity of the variance using a Levene test.

## Acknowledgements

We thank Elisa Pizarro Carbonell, Carlos Barriga Cabanillas, Anja Schmutz, Sandra Gonzalez Sanchez, Lukas Meile, Carlos Federico Ingala, Roman Hüppi, Simon Baumgartner, Benjamin Wilde, Manon Longepierre, Marijn Van de Broek, Leonhard Späth, Inea Lehner, Anna Bugmann, Jianguo Chen, Nicola Haggenmacher and Zita Sartori for their help in the field, and Johan Six for comments on the experimental design. We also thank the Aprisco de Las Corchuelas Field Station and the University of Zurich for the use of their facilities. The study was funded by the Swiss National Science Foundation (PP00P3_170645).

## Author contributions

LS, NE, and CS conceptualised the study; LS and CS designed the experiment; LS, NE, and CS carried out the experiment, LS and CS analysed the data; LS and CS wrote the paper with input from NE.

## Materials & Correspondence

Correspondence and requests for materials should be addressed to Laura Stefan.

## Data availability statement

The data that support the findings of this study are available on Zenodo: https://doi.org/10.5281/zenodo.5223410

## Code availability statement

The R code is available on Zenodo: https://doi.org/10.5281/zenodo.5223410

## Extended Data

**Extended Data Figure 1:**
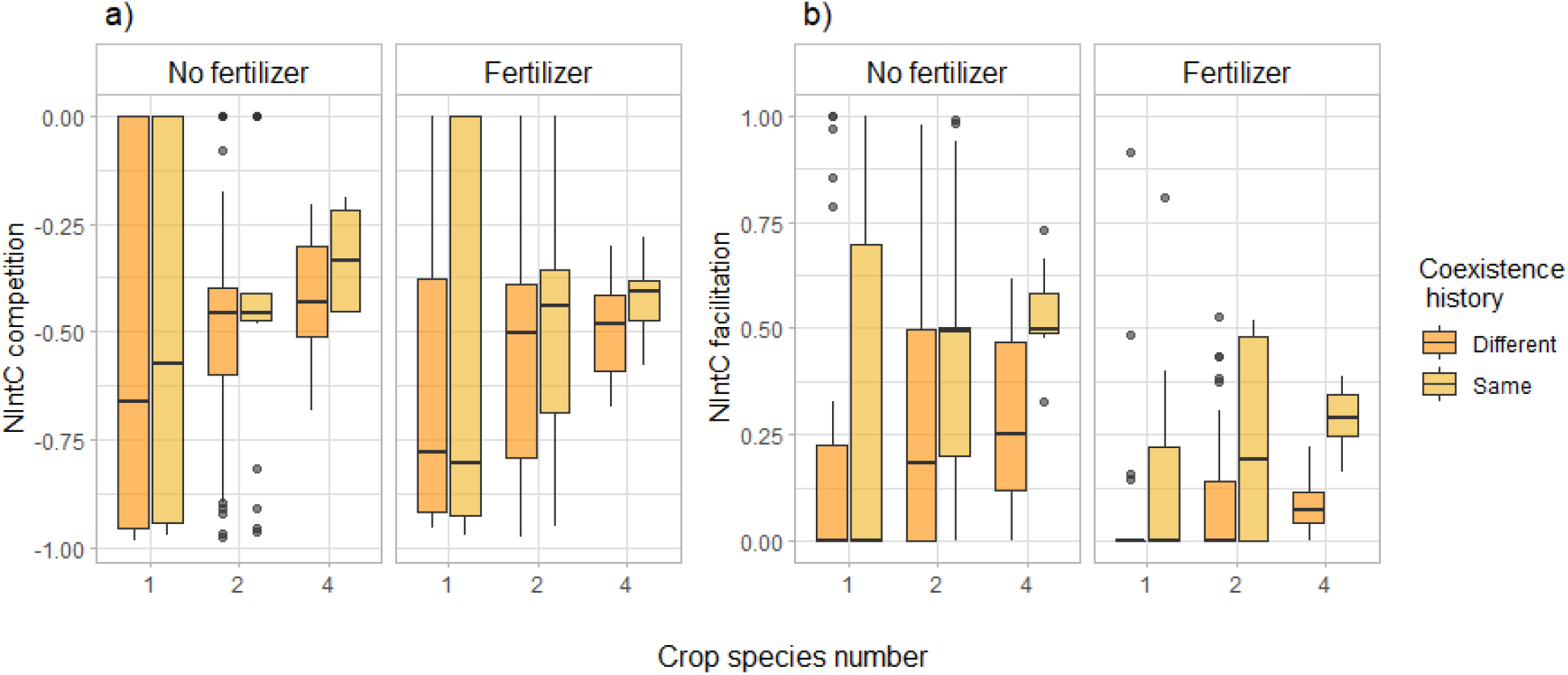
Effects of coexistence history and crop species number on competition (a), and facilitation (b) indexes, for fertilized and unfertilized conditions. “Same coexistence history” indicates that crops were grown in the community their seeds were collected from. “Different coexistence history” refers to crops grown in a community different to the one of their parents. See methods for the index calculations. Horizontal lines represent the median of the data, boxes represent the lower and upper quartiles (25% and 75%), with vertical lines extending from the hinge of the box to the smallest and largest values, no further than 1.5 * the interquartile range. Data beyond the end of the whiskers are outlying and plotted individually. n =276

**Extended Data Table 1.** Type-I Analysis of Variance table of the experimental treatment effects on net, competition and facilitation indexes, in 2020. *DenDF*, degrees of freedom of error term; *NumDF*, degrees of freedom of term; *F-value*, variance ratio; *Pr(>F)*, error probability. P-values in bold are significant at α = 0.05; * (P < 0.05), ** (P < 0.01), *** (P < 0.001). n =276

|  | NumDF | Net |  |  |  | Competition |  |  | Facilitation |  |  |
| --- | --- | --- | --- | --- | --- | --- | --- | --- | --- | --- | --- |
|  |  | DenDF | F value | Pr(>F) |  | DenDF | F value | Pr(>F) | DenDF | F value | Pr(>F) |
| Fertilizer | 1 | 7.02 | 36.269 | **0.0005 \*\*\*** |  | 6.82 | 18.658 | **0.0037 \*\*** | 6.74 | 39.646 | **0.0005 \*\*\*** |
| History | 1 | 238.10 | 31.901 | **4.61E-08 \*\*\*** |  | 232.57 | 9.071 | **0.0029 \*\*** | 236.53 | 38.318 | **2.63E-09 \*\*\*** |
| Monocultures vs. mixtures | 1 | 19.99 | 0.571 | 0.4587 |  | 20.00 | 0.343 | 0.5647 | 19.97 | 0.945 | 0.3425 |
| Diversity | 1 | 19.97 | 0.110 | 0.7438 |  | 20.00 | 0.084 | 0.7756 | 19.90 | 0.159 | 0.6942 |
| Fertilizer x history | 1 | 237.87 | 0.205 | 0.6509 |  | 232.63 | 0.279 | 0.5982 | 236.49 | 0.921 | 0.3383 |
| Fertilizer x mono vs. mix | 1 | 242.37 | 0.695 | 0.4054 |  | 243.37 | 0.036 | 0.8493 | 242.65 | 1.269 | 0.2611 |
| Fertilizer x diversity | 1 | 240.31 | 0.305 | 0.5816 |  | 241.70 | 0.075 | 0.7852 | 240.70 | 0.374 | 0.5415 |
| History x mono vs. mix | 1 | 240.38 | 1.319 | 0.2518 |  | 241.43 | 0.253 | 0.6152 | 240.79 | 2.149 | 0.1440 |
| History x diversity | 1 | 240.50 | 0.912 | 0.3405 |  | 241.93 | 0.279 | 0.5977 | 240.88 | 1.111 | 0.2928 |
| Fertilizer x history x mono vs. mix | 1 | 240.34 | 0.022 | 0.8836 |  | 241.39 | 0.000 | 0.9864 | 240.75 | 0.003 | 0.9567 |
| Fertilizer x history x diversity | 1 | 240.42 | 0.100 | 0.7523 |  | 241.89 | 0.296 | 0.5868 | 240.81 | 0.003 | 0.9542 |

**Extended Data Table 2.**
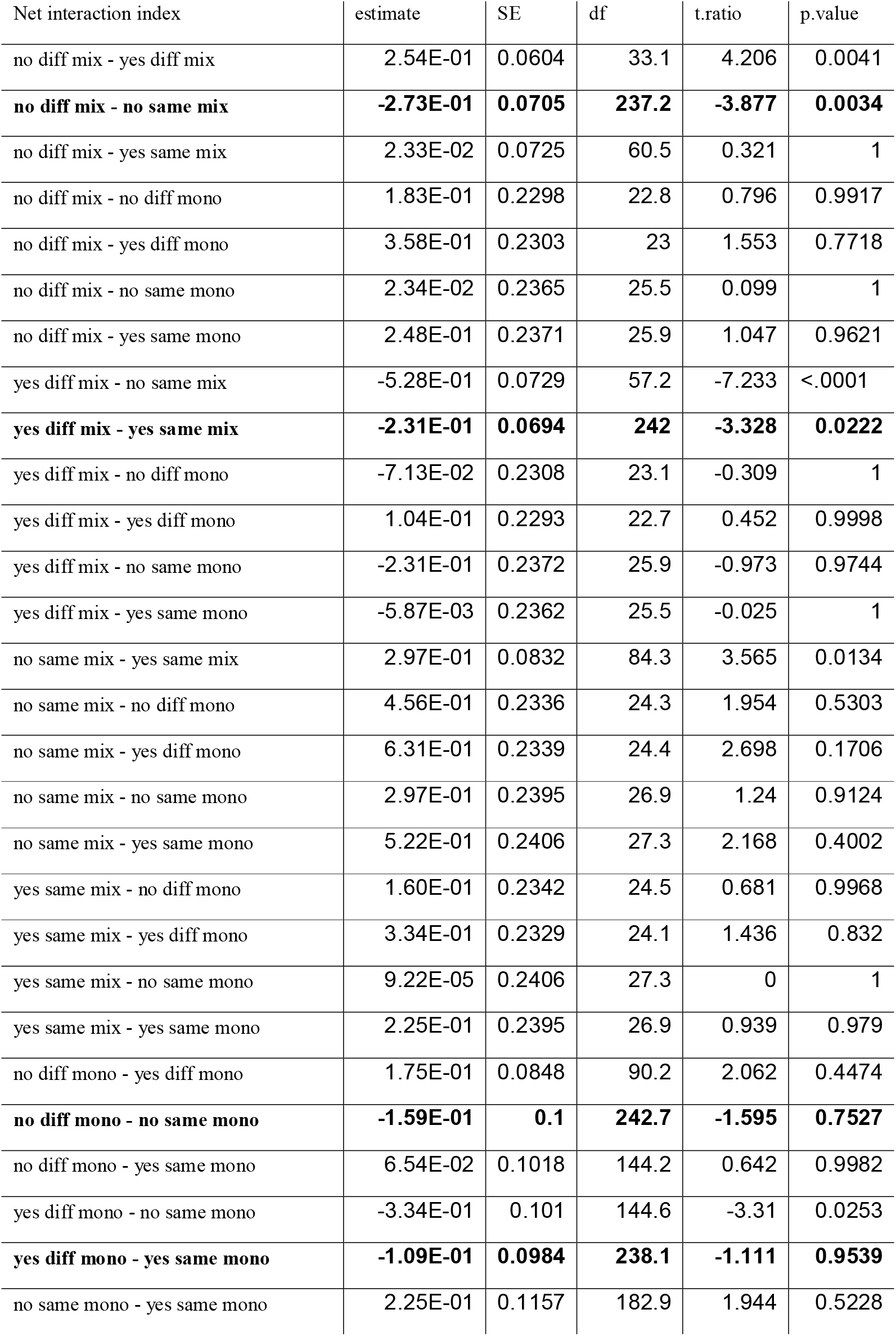
Pairwise comparisons of the effect of net interaction index between fertilizer (yes, no), coexistence history (diff [different], same), and monoculture vs mixture (mix [mixture], mono [monoculture]).

**Extended Data Figure 2:**
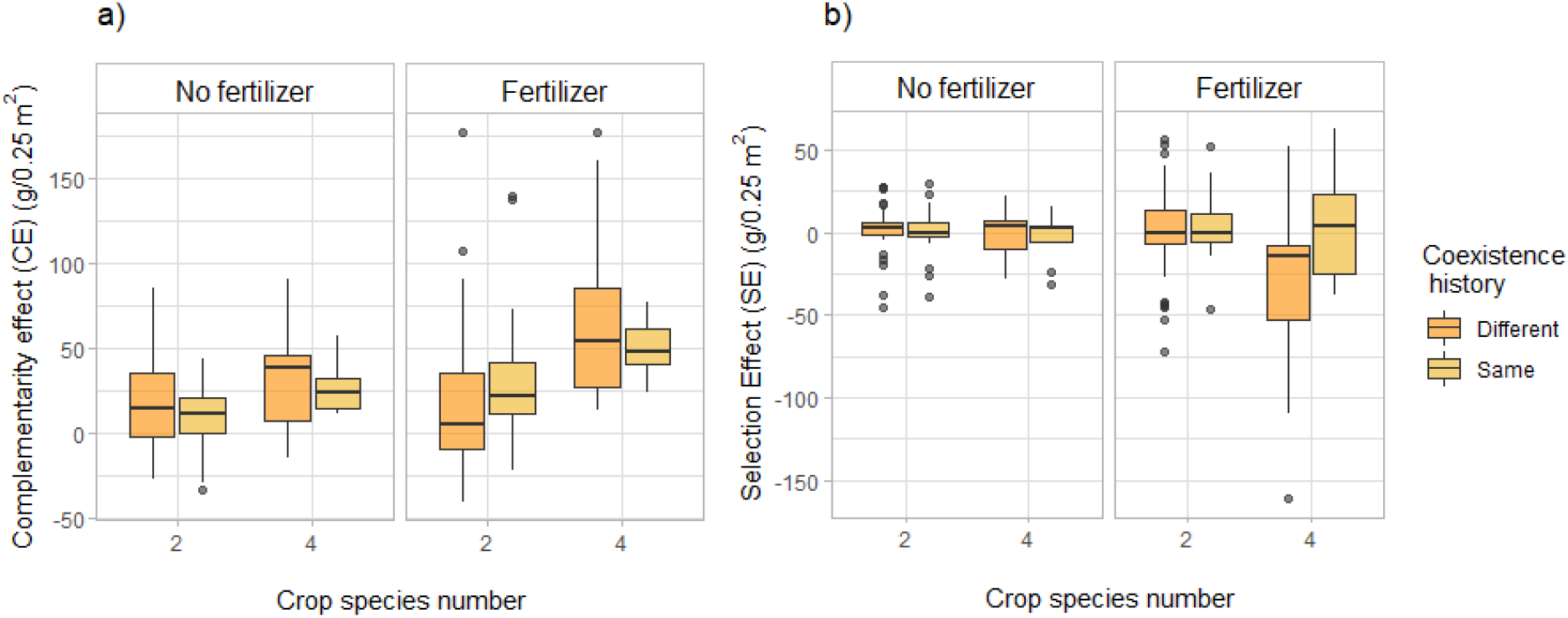
Effects of coexistence history and crop species number on complementarity effect (a) and selection effect (b) in fertilized and unfertilized plots. Horizontal lines represent the median of the data, boxes represent the lower and upper quartiles (25% and 75%), with vertical lines extending from the hinge of the box to the smallest and largest values, no further than 1.5 * the interquartile range. Data beyond the end of the whiskers are outlying and plotted individually. n=204

**Extended Data Table 3.** Type-I Analysis of Variance table of the experimental treatment effects on net, complementarity, and selection effects in 2020. *DenDF*, degrees of freedom of error term; *NumDF*, degrees of freedom of term; *F-value*, variance ratio; *Pr(>F)*, error probability. P-values in bold are significant at α = 0.1;. (P < 0.1); * (P < 0.05), ** (P < 0.01), *** (P < 0.001). n=204

|  | *NumDF* | *Net effect* |  |  | *Complementarity effect* |  |  | *Selection effect* |  |  |
| --- | --- | --- | --- | --- | --- | --- | --- | --- | --- | --- |
|  |  | *DenDF* | *F value* | *Pr(>F)* | *DenDF* | *F value* | *Pr(>F)* | *DenDF* | *F value* | *Pr(>F)* |
| *Fertilizer* | 1 | 7.64 | 1.005 | 0.3468 | 7.38 | 2.684 | 0.14312 | 7.69 | 1.295 | 0.2894 |
| *History* | 1 | 177.34 | 0.162 | 0.6876 | 179.89 | 1.567 | 0.2123 | 178.62 | 1.512 | 0.2204 |
| *Diversity* | 1 | 14.90 | 2.583 | 0.1290 | 14.87 | 5.887 | **0.0285** * | 15.06 | 4.009 | **0.0636** . |
| *Fertilizer x history* | 1 | 177.08 | 9.595 | **0.0023** ** | 179.80 | 2.719 | 0.1009 | 178.66 | 2.498 | 0.1158 |
| *Fertilizer x diversity* | 1 | 173.48 | 0.197 | 0.6581 | 174.92 | 5.579 | **0.0193** * | 178.22 | 6.092 | **0.0145** * |
| *History x diversity* | 1 | 173.66 | 0.052 | 0.8191 | 175.06 | 1.399 | 0.2385 | 178.42 | 3.165 | **0.0769** . |
| *Fertilizer x history x diversity* | 1 | 173.36 | 0.046 | 0.8297 | 174.85 | 2.026 | 0.1564 | 178.35 | 4.093 | **0.0446** * |

**Extended Data Table 4.**
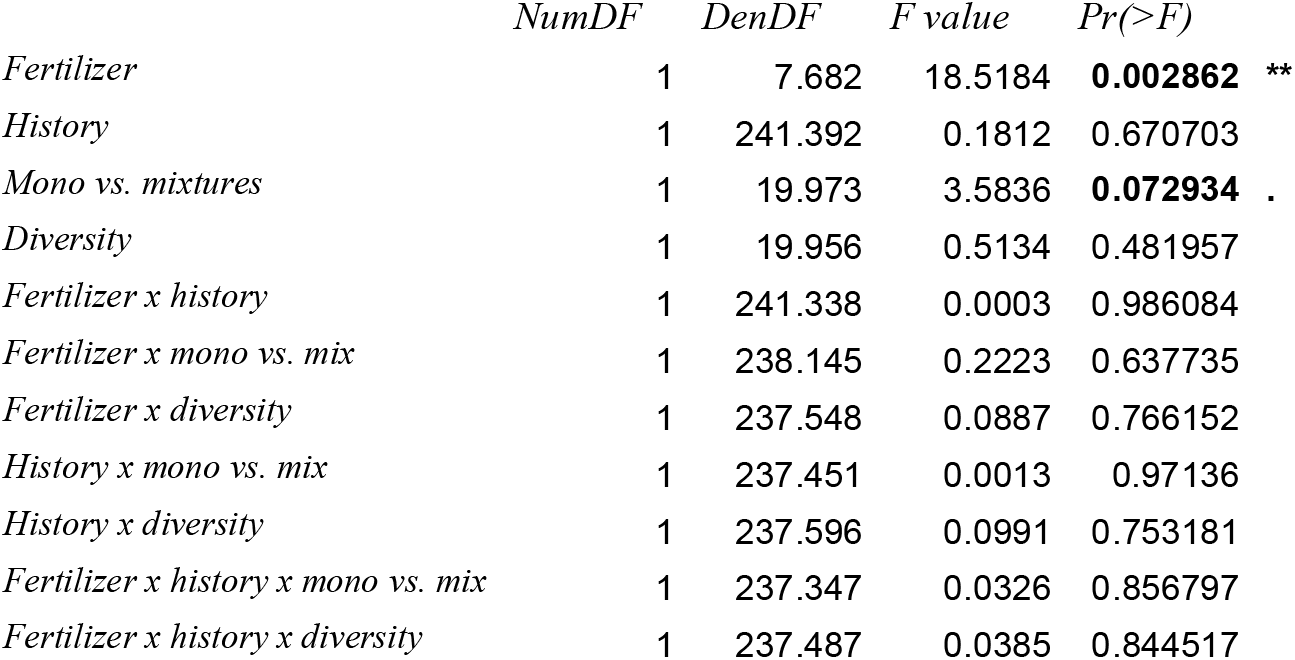
Type-I Analysis of Variance table of the experimental treatment effects on total crop yield per plot (square-root transformed) *DenDF*, degrees of freedom of error term; *NumDF*, degrees of freedom of term; *F-value*, variance ratio; *Pr(>F)*, error probability. P-values in bold are significant at α = 0.1;. (P < 0.1); * (P < 0.05), ** (P < 0.01), *** (P < 0.001). n=276

**Extended Data Figure 3:**
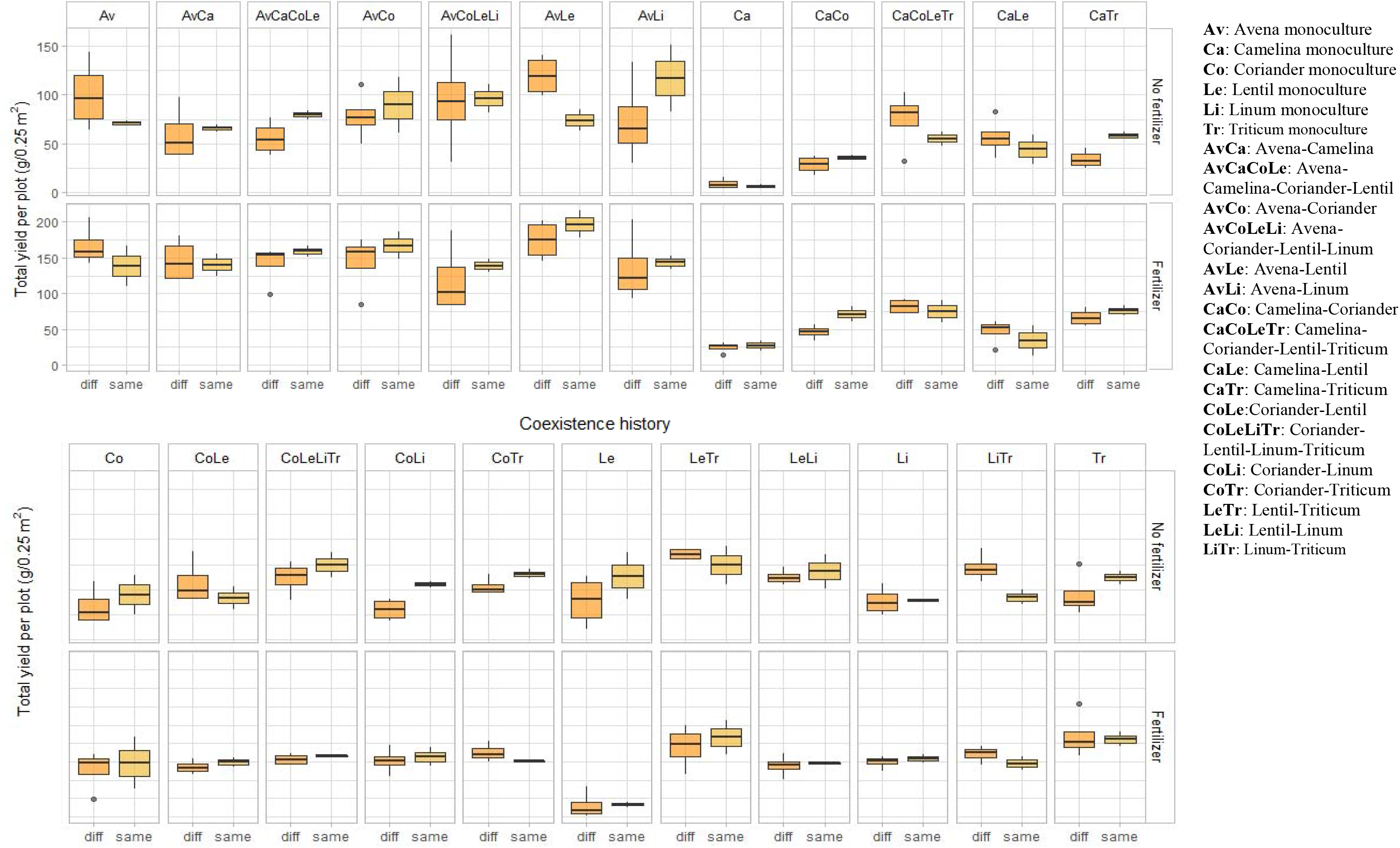
Effects of coexistence history of total yield per plot, per species combination. “Same coexistence history” indicates that crops were grown in the community their seeds were collected from. “Different coexistence history” refers to crops grown in a community different to the one of their parents.

**Extended Data Figure 5:**
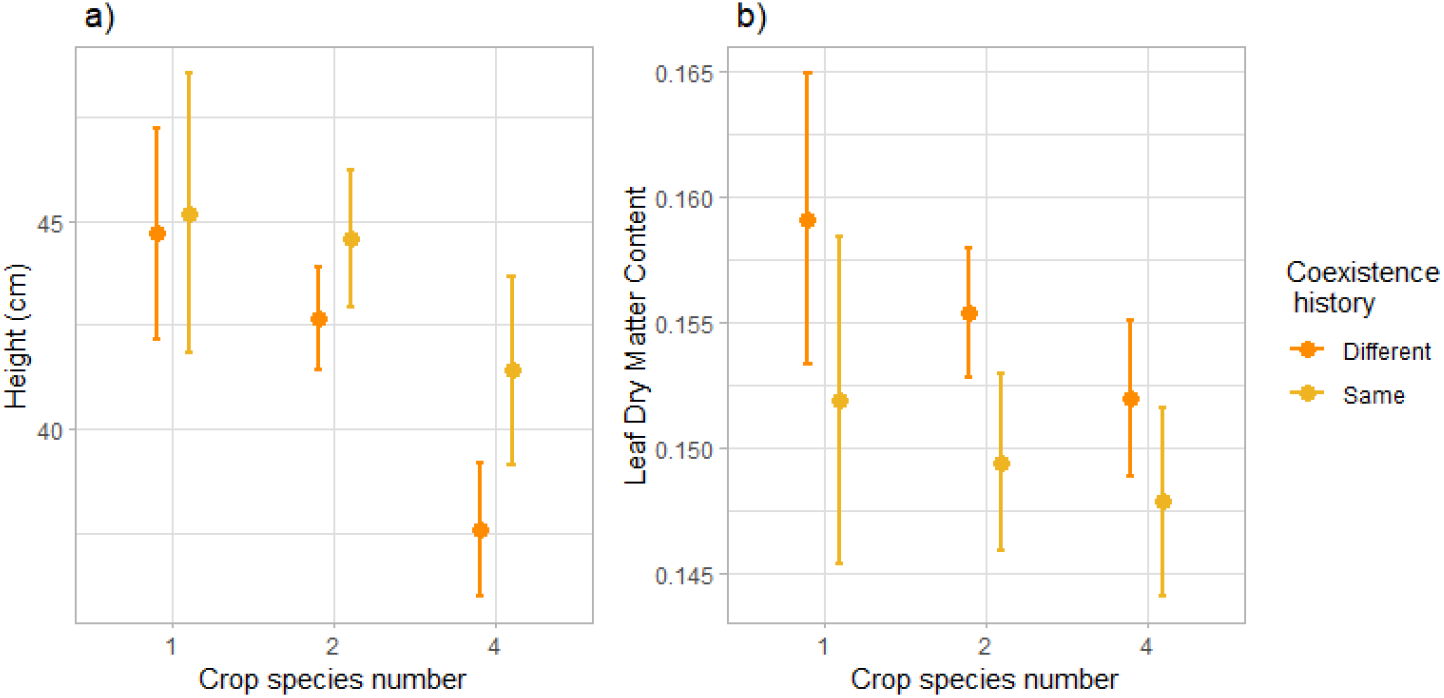
Effects of coexistence history and crop species number on mean height (in cm) (a) and LDMC (b). Dots represent the averaged values across species and plots; lines represent the standard error. n =1726

**Extended Data Figure 6:**
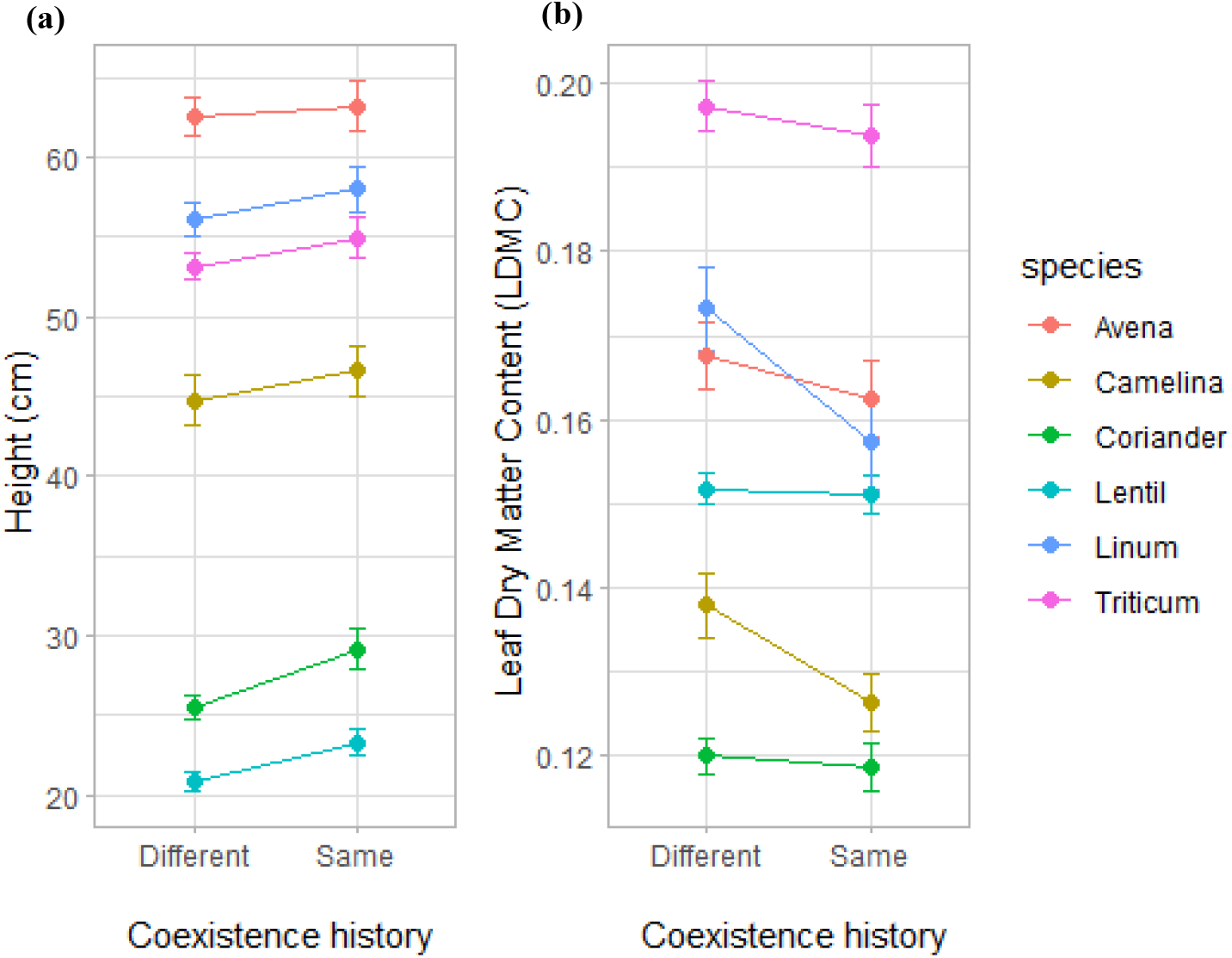
Mean height (cm) (a) and LDMC (b) according to their coexistence history, for the six species considered in our study. Dots represent the averaged values across species and plots; lines represent the standard error. n =1726

**Extended Data Table 5.** Type-I Analysis of Variance table of the experimental treatment effects on mean and coefficient of variation of height, per species per plot (species level) *DenDF*, degrees of freedom of error term; *NumDF*, degrees of freedom of term; *F-value*, variance ratio; *Pr(>F)*, error probability. P-values in bold are significant at α = 0.05; * (P < 0.05), ** (P < 0.01), *** (P < 0.001), n=1726

|  | Mean |  |  |  | Coefficient of variation |  |  |
| --- | --- | --- | --- | --- | --- | --- | --- |
|  | *NumDF* | *DenDF* | *F value* | *Pr(>F)* | *DenDF* | *F value* | *Pr(>F)* |
| *Fertilizer* | 1 | 7.9 | 0.2292 | 0.645078 | 7.04 | 16.8065 | **4.51E-03** ** |
| *History* | 1 | 169.16 | 4.2929 | **0.039789** * | 197.26 | 0.077 | 0.781684 |
| *Mono vs. mixtures* | 1 | 23.57 | 0.1617 | 0.691188 | 45.01 | 0.325 | 0.571472 |
| *Diversity* | 1 | 10.68 | 0.121 | 0.73467 | 45.48 | 0.2586 | 0.61356 |
| *Fertilizer x history* | 1 | 168.93 | 0.1986 | 0.656442 | 197.18 | 5.8068 | **0.016883** * |
| *Fertilizer x mono vs. mix* | 1 | 416.7 | 9.2129 | **0.002554** ** | 487.47 | 5.0102 | **0.025648** * |
| *Fertilizer x diversity* | 1 | 118.78 | 0.0013 | 0.971368 | 127.23 | 0.0009 | 0.976698 |
| *History x mono vs. mix* | 1 | 418.29 | 1.435 | 0.231632 | 483.31 | 0.2095 | 0.647359 |
| *History x diversity* | 1 | 119.33 | 0.6601 | 0.418149 | 128.24 | 0.1385 | 0.710429 |
| *Fertilizer x history x mono vs. mix* | 1 | 417.97 | 0.0046 | 0.945955 | 483.3 | 0.5825 | 0.445694 |
| *Fertilizer x history x diversity* | 1 | 119.15 | 0.065 | 0.79923 | 128.25 | 0.0665 | 0.796946 |

**Extended Data Table 6.** Type-I Analysis of Variance table of the experimental treatment effects on mean and coefficient of variation of width, per species per plot (species level) *DenDF*, degrees of freedom of error term; *NumDF*, degrees of freedom of term; *F-value*, variance ratio; *Pr(>F)*, error probability. P-values in bold are significant at α = 0.1;. (P < 0.1); * (P < 0.05), ** (P < 0.01), *** (P < 0.001), n=1726

|  | Mean |  |  |  | Coefficient of variation |  |  |
| --- | --- | --- | --- | --- | --- | --- | --- |
|  | *NumDF* | *DenDF* | *F value* | *Pr(>F)* | *DenDF* | *F value* | *Pr(>F)* |
| *Fertilizer* | 1 | 7.59 | 8.5571 | **0.02027** * | 7.34 | 4.8793 | **6.12E-02** |
| *History* | 1 | 531.5 | 2.0724 | 0.15057 | 233.05 | 2.5024 | 0.11503 |
| *Mono vs. mixtures* | 1 | 19.68 | 0.9369 | 0.34482 | 534.07 | 0.0141 | 0.90552 |
| *Diversity* | 1 | 10.47 | 0.0639 | 0.80529 | 144.7 | 0.8817 | 0.3493 |
| *Fertilizer x history* | 1 | 530.95 | 1.6511 | 0.19937 | 232.84 | 0.0033 | 0.9541 |
| *Fertilizer x mono vs. mix* | 1 | 523.5 | 2.5201 | 0.11301 | 533.17 | 0.7796 | 0.37767 |
| *Fertilizer x diversity* | 1 | 526.73 | 3.905 | **0.04866** * | 142.23 | 0.2794 | 0.59791 |
| *History x mono vs. mix* | 1 | 522.6 | 0.1858 | 0.66657 | 528.21 | 1.4807 | 0.2242 |
| *History x diversity* | 1 | 526.98 | 1.1295 | 0.28837 | 144.44 | 1.1028 | 0.2954 |
| *Fertilizer x history x mono vs. mix* | 1 | 522.34 | 0.054 | 0.81631 | 528.23 | 1.7197 | 0.1903 |
| *Fertilizer x history x diversity* | 1 | 526.26 | 0.9072 | 0.3413 | 144.54 | 0.8736 | 0.35153 |

**Extended Data Table 7.**
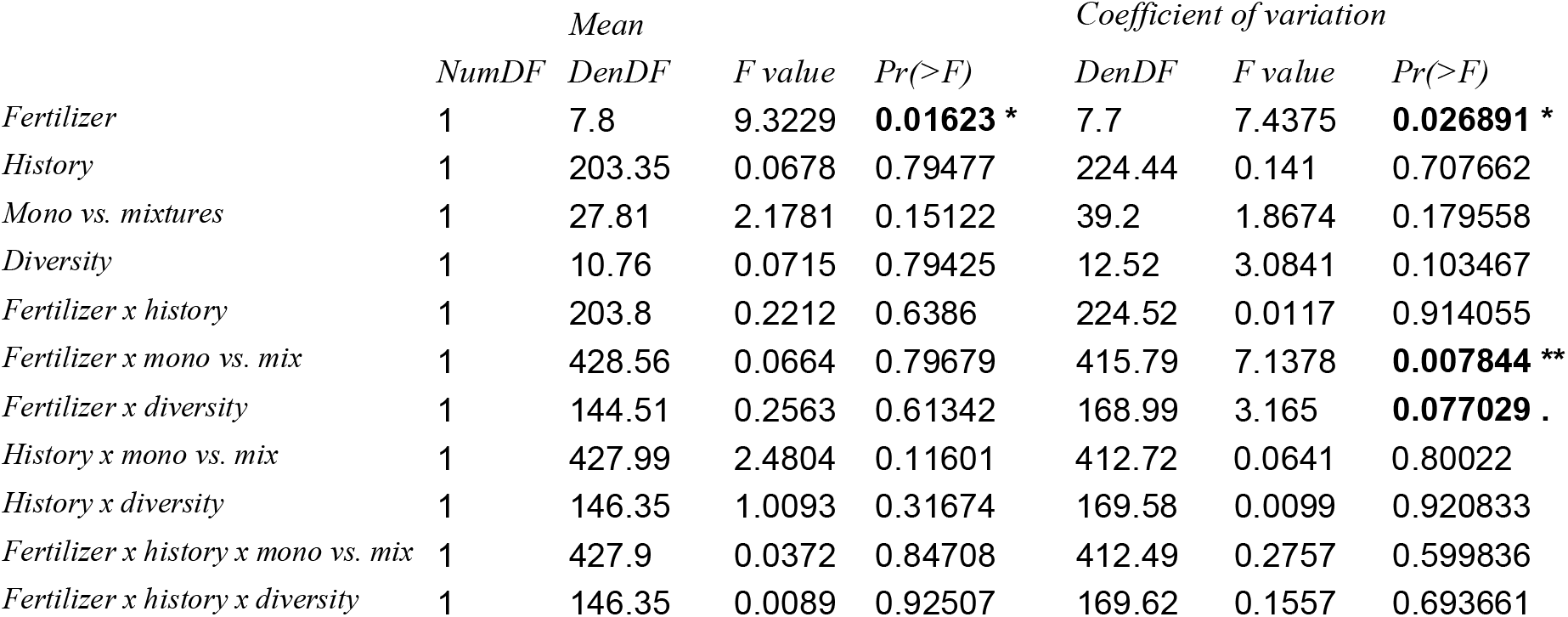
Type-I Analysis of Variance table of the experimental treatment effects on mean and coefficient of variation of SLA, per species per plot (species level) *DenDF*, degrees of freedom of error term; *NumDF*, degrees of freedom of term; *F-value*, variance ratio; *Pr(>F)*, error probability. P-values in bold are significant at α = 0.1;. (P < 0.1); * (P < 0.05), ** (P < 0.01), *** (P < 0.001), n=1726

**Extended Data Table 8.** Type-I Analysis of Variance table of the experimental treatment effects on mean and coefficient of variation of LDMC, per species per plot (species level) *DenDF*, degrees of freedom of error term; *NumDF*, degrees of freedom of term; *F-value*, variance ratio; *Pr(>F)*, error probability. P-values in bold are significant at α = 0.1;. (P < 0.1); * (P < 0.05), ** (P < 0.01), *** (P < 0.001), n=1726

|  | Mean |  |  |  | Coefficient of variation |  |  |
| --- | --- | --- | --- | --- | --- | --- | --- |
|  | *NumDF* | *DenDF* | *F value* | *Pr(>F)* | *DenDF* | *F value* | *Pr(>F)* |
| *Fertilizer* | 1 | 7.86 | 8.0352 | **0.02239** * | 186.4 | 12.783 | **4.46E-04** *** |
| *History* | 1 | 183.84 | 3.5956 | **0.05950** . | 190.46 | 0.0447 | 0.832786 |
| *Mono vs. mixtures* | 1 | 19.67 | 0.1589 | 0.69442 | 516.86 | 0.0353 | 0.851015 |
| *Diversity* | 1 | 5.24 | 0.1083 | 0.75481 | 116.92 | 0.6369 | 0.426449 |
| *Fertilizer x history* | 1 | 181.59 | 0.0432 | 0.83566 | 190.48 | 3.1314 | **0.078397** . |
| *Fertilizer x mono vs. mix* | 1 | 467.36 | 0.0294 | 0.86399 | 515.71 | 0.6107 | 0.434889 |
| *Fertilizer x diversity* | 1 | 115.57 | 0.3376 | 0.56232 | 114.6 | 3.8577 | **0.051939** . |
| *History x mono vs. mix* | 1 | 468.83 | 0.0043 | 0.94775 | 515.4 | 0.2779 | 0.598296 |
| *History x diversity* | 1 | 116.59 | 1.0418 | 0.30953 | 114.82 | 0.9337 | 0.335923 |
| *Fertilizer x history x mono vs. mix* | 1 | 468.38 | 0.0166 | 0.89752 | 515.36 | 1.6235 | 0.203176 |
| *Fertilizer x history x diversity* | 1 | 116.48 | 0.0016 | 0.96772 | 114.88 | 1.3649 | 0.24511 |

**Extended Data Table 9.** Type-I Analysis of Variance table of the experimental treatment effects on mean and coefficient of variation of mass per seed, per species per plot (species level) *DenDF*, degrees of freedom of error term; *NumDF*, degrees of freedom of term; *F-value*, variance ratio; *Pr(>F)*, error probability. P-values in bold are significant at α = 0.1;. (P < 0.1); * (P < 0.05), ** (P < 0.01), *** (P < 0.001), n=1726

|  | *Mean* |  |  |  | *Coefficient of variation* |  |  |
| --- | --- | --- | --- | --- | --- | --- | --- |
|  | *NumDF* | *DenDF* | *F value* | *Pr(>F)* | *DenDF* | *F value* | *Pr(>F)* |
| *Fertilizer* | 1 | 210.65 | 4.3651 | **0.03788** * | 180.24 | 0.1253 | 7.24E-01 |
| *History* | 1 | 211.89 | 0.0491 | 0.82487 | 182.21 | 0.0853 | 0.770551 |
| *Mono vs. mixtures* | 1 | 20.08 | 10.0297 | **0.00483** ** | 33.75 | 17.0854 | **0.000223** *** |
| *Diversity* | 1 | 10.77 | 1.3367 | 0.27261 | 8.88 | 0.1203 | 0.736758 |
| *Fertilizer x history* | 1 | 211.88 | 0.6633 | 0.4163 | 182.12 | 1.6601 | 0.199224 |
| *Fertilizer x mono vs. mix* | 1 | 493.44 | 0.602 | 0.43818 | 474.87 | 1.16 | 0.282008 |
| *Fertilizer x diversity* | 1 | 137.39 | 0.3993 | 0.52848 | 118.13 | 1.4557 | 0.230024 |
| *History x mono vs. mix* | 1 | 493.59 | 1.4337 | 0.23174 | 470.19 | 4.9519 | **0.026536** * |
| *History x diversity* | 1 | 138.08 | 0.2159 | 0.64289 | 119.33 | 0.2756 | 0.600592 |
| *Fertilizer x history x mono vs. mix* | 1 | 493.59 | 2.7072 | 0.10053 | 470.29 | 0.0587 | 0.808609 |
| *Fertilizer x history x diversity* | 1 | 138.1 | 0.6308 | 0.42842 | 119.29 | 0.0329 | 0.856454 |

**Extended Data Table 10.**
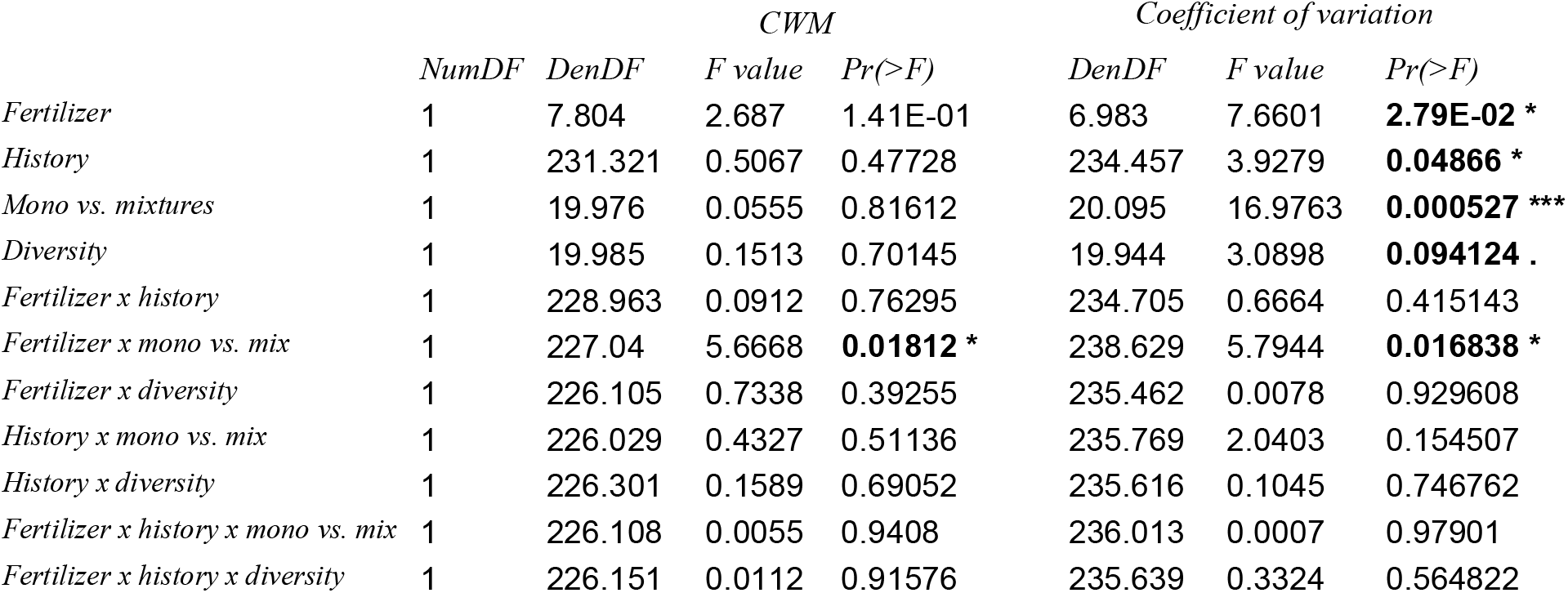
Type-I Analysis of Variance table of the experimental treatment effects on community-weighted mean and coefficient of variation of height, per plot (community level) *DenDF*, degrees of freedom of error term; *NumDF*, degrees of freedom of term; *F-value*, variance ratio; *Pr(>F)*, error probability. P-values in bold are significant at α = 0.1;. (P < 0.1); * (P < 0.05), ** (P < 0.01), *** (P < 0.001). n=271

|  | *NumDF* | *DenDF* | *CWM* |  | *Coefficient of variation* |  |  |
| --- | --- | --- | --- | --- | --- | --- | --- |
|  |  |  | *F value* | *Pr(>F)* | *DenDF* | *F value* | *Pr(>F)* |
| *Fertilizer* | 1 | 7.804 | 2.687 | 1.41E-01 | 6.983 | 7.6601 | **2.79E-02** * |
| *History* | 1 | 231.321 | 0.5067 | 0.47728 | 234.457 | 3.9279 | **0.04866** * |
| *Mono vs. mixtures* | 1 | 19.976 | 0.0555 | 0.81612 | 20.095 | 16.9763 | **0.000527** *** |
| *Diversity* | 1 | 19.985 | 0.1513 | 0.70145 | 19.944 | 3.0898 | **0.094124** . |
| *Fertilizer x history* | 1 | 228.963 | 0.0912 | 0.76295 | 234.705 | 0.6664 | 0.415143 |
| *Fertilizer x mono vs. mix* | 1 | 227.04 | 5.6668 | **0.01812** * | 238.629 | 5.7944 | **0.016838** * |
| *Fertilizer x diversity* | 1 | 226.105 | 0.7338 | 0.39255 | 235.462 | 0.0078 | 0.929608 |
| *History x mono vs. mix* | 1 | 226.029 | 0.4327 | 0.51136 | 235.769 | 2.0403 | 0.154507 |
| *History x diversity* | 1 | 226.301 | 0.1589 | 0.69052 | 235.616 | 0.1045 | 0.746762 |
| *Fertilizer x history x mono vs. mix* | 1 | 226.108 | 0.0055 | 0.9408 | 236.013 | 0.0007 | 0.97901 |
| *Fertilizer x history x diversity* | 1 | 226.151 | 0.0112 | 0.91576 | 235.639 | 0.3324 | 0.564822 |

**Extended Data Table 11.** Type-I Analysis of Variance table of the experimental treatment effects on community-weighted mean and coefficient of variation of width, per plot (community level) *DenDF*, degrees of freedom of error term; *NumDF*, degrees of freedom of term; *F-value*, variance ratio; *Pr(>F)*, error probability. P-values in bold are significant at α = 0.1;. (P < 0.1); * (P < 0.05), ** (P < 0.01), *** (P < 0.001), n =271

|  | *NumDF* | *DenDF* | *CWM* |  | *Coefficient of variation* |  |  |
| --- | --- | --- | --- | --- | --- | --- | --- |
|  |  |  | *F value* | *Pr(>F)* | *DenDF* | *F value* | *Pr(>F)* |
| *Fertilizer* | 1 | 7.484 | 6.0869 | **4.09E-02** * | 7.352 | 10.7862 | **1.25E-02** * |
| *History* | 1 | 233.262 | 0.001 | 0.974539 | 239.502 | 0.2976 | 0.585869 |
| *Mono vs. mixtures* | 1 | 19.917 | 1.7012 | 0.207003 | 20.153 | 15.6397 | **0.000773** *** |
| *Diversity* | 1 | 19.935 | 0.383 | 0.543019 | 19.819 | 2.4097 | 0.13641 |
| *Fertilizer x history* | 1 | 231.77 | 1.6122 | 0.205462 | 239.1 | 0.0345 | 0.852701 |
| *Fertilizer x mono vs. mix* | 1 | 228.97 | 11.0067 | **0.001056** ** | 237.11 | 8.7687 | **0.003376** ** |
| *Fertilizer x diversity* | 1 | 227.323 | 3.9302 | **0.048631** * | 233.942 | 1.1993 | 0.274588 |
| *History x mono vs. mix* | 1 | 227.213 | 0.1337 | 0.714952 | 234.231 | 4.7012 | **0.031149** * |
| *History x diversity* | 1 | 227.671 | 0.1338 | 0.714862 | 234.017 | 0.029 | 0.864848 |
| *Fertilizer x history x mono vs. mix* | 1 | 227.362 | 0.391 | 0.532421 | 234.403 | 2.0867 | 0.149924 |
| *Fertilizer x history x diversity* | 1 | 227.418 | 0.3123 | 0.576817 | 233.993 | 0.6311 | 0.427744 |

**Extended Data Table 12.**
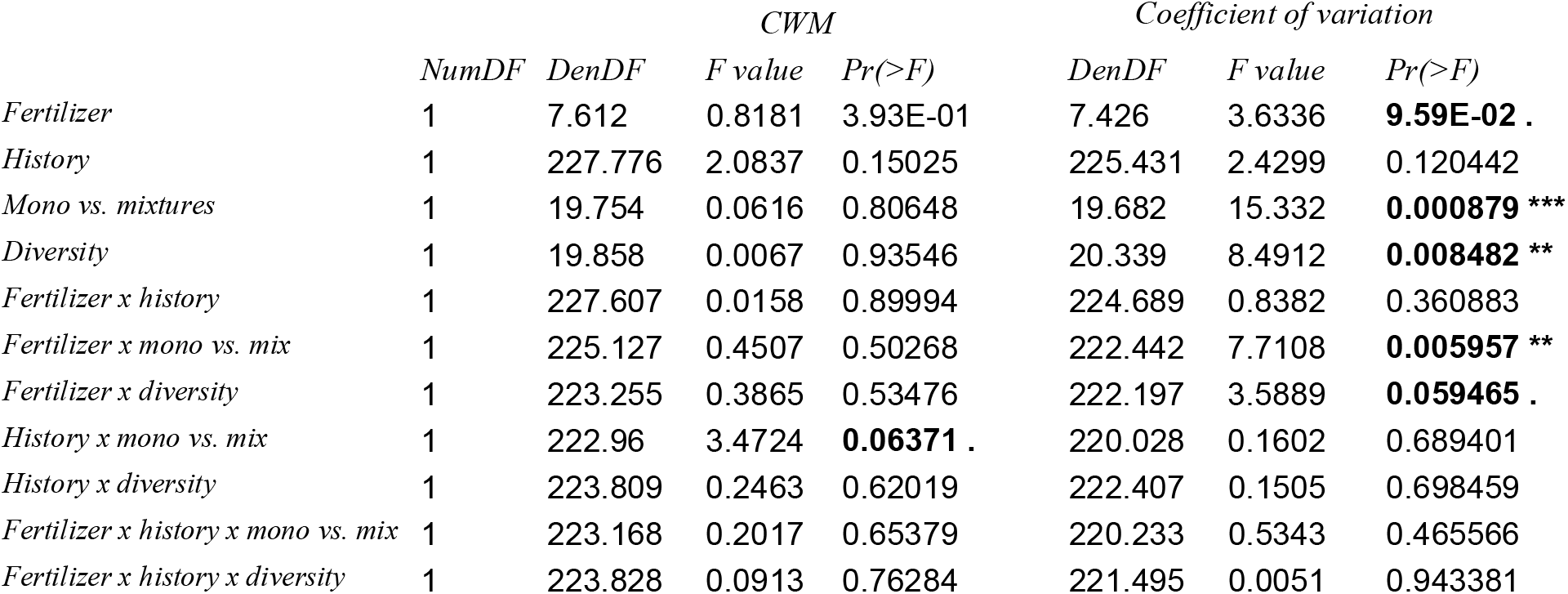
Type-I Analysis of Variance table of the experimental treatment effects on community-weighted mean and coefficient of variation of SLA, per plot (community level) *DenDF*, degrees of freedom of error term; *NumDF*, degrees of freedom of term; *F-value*, variance ratio; *Pr(>F)*, error probability. P-values in bold are significant at α = 0.1;. (P < 0.1); * (P < 0.05), ** (P < 0.01), *** (P < 0.001), n=271

**Extended Data Table 13.** Type-I Analysis of Variance table of the experimental treatment effects on community-weighted mean and coefficient of variation of LDMC, per plot (community level) *DenDF*, degrees of freedom of error term; *NumDF*, degrees of freedom of term; *F-value*, variance ratio; *Pr(>F)*, error probability. P-values in bold are significant at α = 0.1;. (P < 0.1); * (P < 0.05), ** (P < 0.01), *** (P < 0.001), n=271

|  | *CWM* |  |  |  | *Coefficient of variation* |  |  |
| --- | --- | --- | --- | --- | --- | --- | --- |
|  | *NumDF* | *DenDF* | *F value* | *Pr(>F)* | *DenDF* | *F value* | *Pr(>F)* |
| *Fertilizer* | 1 | 7.809 | 10.5893 | **1.20E-02** * | 7.368 | 2.1001 | 1.88E-01 |
| *History* | 1 | 225.249 | 4.3323 | **0.03853** * | 233.338 | 4.1789 | **0.042053** * |
| *Mono vs. mixtures* | 1 | 19.998 | 0.0614 | 0.80688 | 20.123 | 10.747 | **0.003737** ** |
| *Diversity* | 1 | 20.012 | 0.0293 | 0.86579 | 19.46 | 0.7439 | 0.39891 |
| *Fertilizer x history* | 1 | 223.935 | 0.598 | 0.44016 | 233.38 | 0.9977 | 0.318908 |
| *Fertilizer x mono vs. mix* | 1 | 222.756 | 0.3654 | 0.54615 | 236.325 | 4.6325 | **0.032385** * |
| *Fertilizer x diversity* | 1 | 221.841 | 0.0235 | 0.87843 | 232.776 | 0.0154 | 0.901256 |
| *History x mono vs. mix* | 1 | 221.728 | 0.1477 | 0.70108 | 233.192 | 0.4468 | 0.504525 |
| *History x diversity* | 1 | 222.051 | 0.0037 | 0.95178 | 233 | 0.3198 | 0.572283 |
| *Fertilizer x history x mono vs. mix* | 1 | 221.903 | 0.1505 | 0.69841 | 233.579 | 2.708 | 0.101189 |
| *Fertilizer x history x diversity* | 1 | 221.981 | 0.3306 | 0.56587 | 233.206 | 0.8702 | 0.351866 |

**Extended Data Table 14.** Type-I Analysis of Variance table of the experimental treatment effects on community-weighted mean and coefficient of variation of mass per seed, per plot (community level) *DenDF*, degrees of freedom of error term; *NumDF*, degrees of freedom of term; *F-value*, variance ratio; *Pr(>F)*, error probability. P-values in bold are significant at α = 0.1;. (P < 0.1); * (P < 0.05), ** (P < 0.01), *** (P < 0.001), n=271

|  | *CWM* |  |  |  | *Coefficient of variation* |  |  |
| --- | --- | --- | --- | --- | --- | --- | --- |
|  | *NumDF* | *DenDF* | *F value* | *Pr(>F)* | *DenDF* | *F value* | *Pr(>F)* |
| *Fertilizer* | 1 | 7.319 | 2.4734 | 0.15792 | 6.612 | 1.0304 | 3.46E-01 |
| *History* | 1 | 240.762 | 0.3199 | 0.57217 | 225.443 | 1.4213 | 0.234444 |
| *Mono vs. mixtures* | 1 | 20.012 | 0.1269 | 0.72535 | 20.021 | 14.3694 | **0.001145** ** |
| *Diversity* | 1 | 19.996 | 0.004 | 0.9499 | 19.95 | 0.5302 | 0.474995 |
| *Fertilizer x history* | 1 | 235.451 | 0.0844 | 0.77174 | 226.003 | 0.0292 | 0.864583 |
| *Fertilizer x mono vs. mix* | 1 | 238.14 | 0.9245 | 0.33727 | 239.091 | 0.1291 | 0.719721 |
| *Fertilizer x diversity* | 1 | 235.395 | 0.3647 | 0.54651 | 236.191 | 0.2304 | 0.631662 |
| *History x mono vs. mix* | 1 | 235.632 | 1.5019 | 0.22161 | 236.507 | 5.4785 | **0.020084** * |
| *History x diversity* | 1 | 235.537 | 0.0009 | 0.9767 | 236.403 | 0.1301 | 0.718673 |
| *Fertilizer x history x mono vs. mix* | 1 | 235.858 | 3.5455 | **0.06094** . | 236.836 | 0.3515 | 0.553841 |
| *Fertilizer x history x diversity* | 1 | 235.548 | 0.3137 | 0.57596 | 236.483 | 0.1133 | 0.736717 |

**Extended Data Figure 7:**
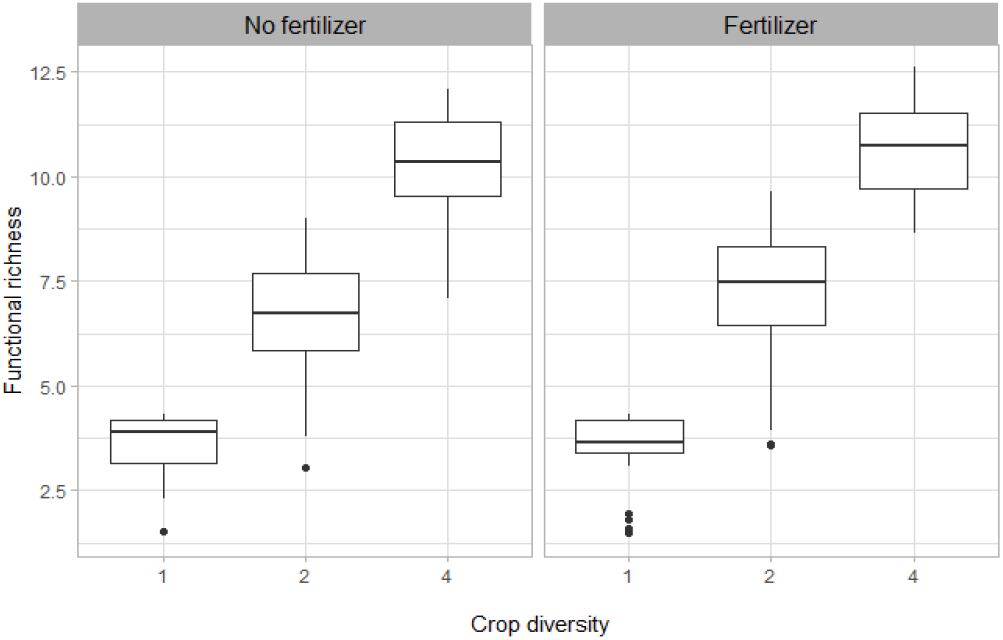
Functional richness in response to crop species diversity, in fertilized and unfertilized plots. N=271

**Extended Data Table 15.**
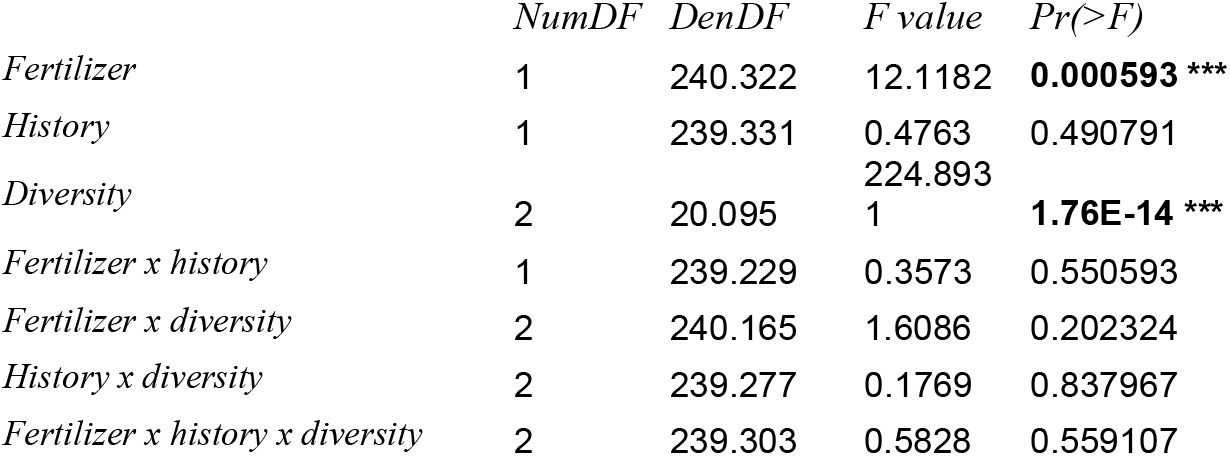
Type-I Analysis of Variance table of the experimental treatment effects on functional richness. *DenDF*, degrees of freedom of error term; *NumDF*, degrees of freedom of term; *F-value*, variance ratio; *Pr(>F)*, error probability. P-values in bold are significant at α = 0.1;. (P < 0.1); * (P < 0.05), ** (P < 0.01), *** (P < 0.001). n=271

**Extended Data Figure 8:**
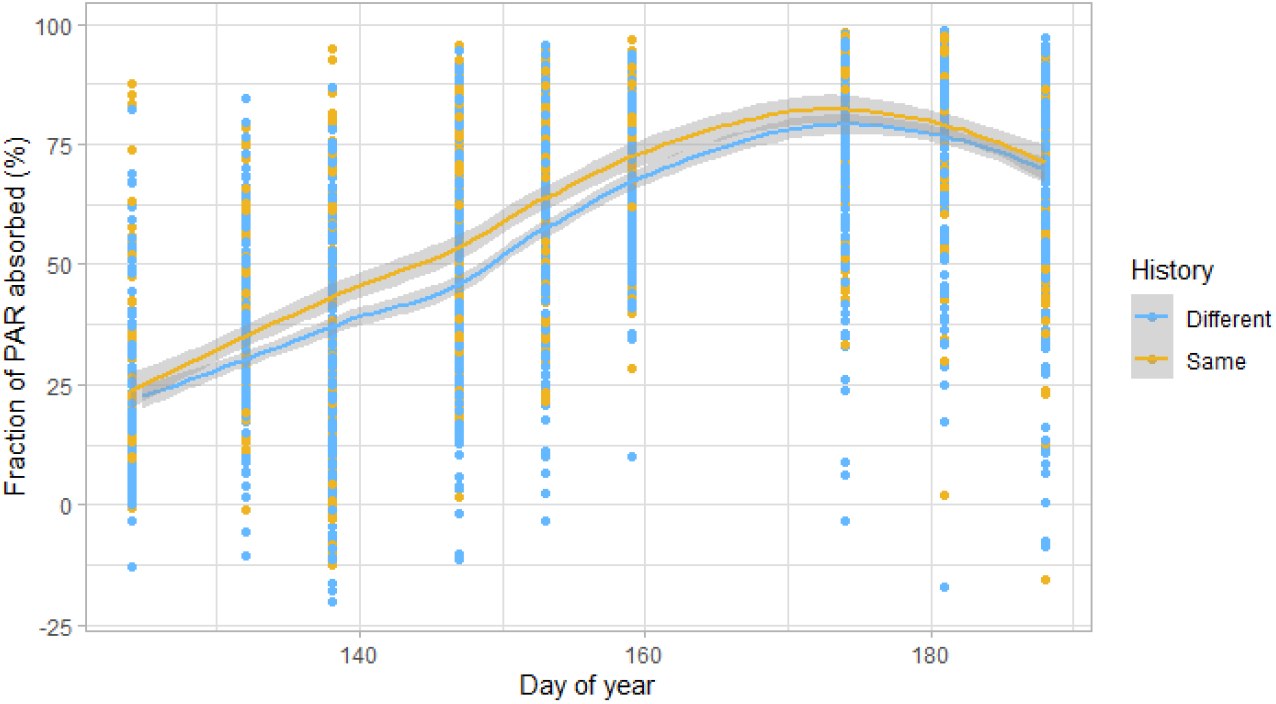
Fraction of PAR absorbed (in %) according to the day of year, for plants with the same or different coexistence history. The lines represent local polynomial regression fittings, with the grey area representing the 0.95 confidence interval. n=2484.

**Extended Data Table 16.**
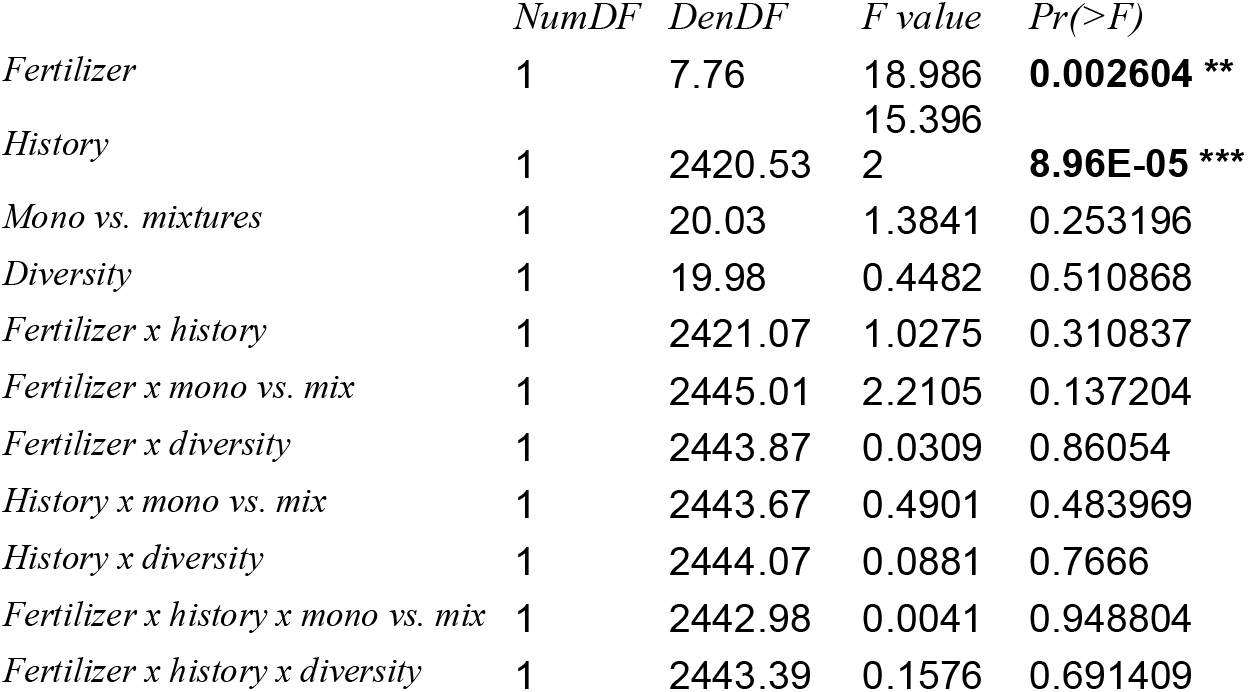
Type-I Analysis of Variance table of the experimental treatment effects on FPAR. *DenDF*, degrees of freedom of error term; *NumDF*, degrees of freedom of term; *F-value*, variance ratio; *Pr(>F)*, error probability. P-values in bold are significant at α = 0.05; * (P < 0.05), ** (P < 0.01), *** (P < 0.001). n=2484

